# GABAergic interneurons contribute to the fatal seizure phenotype of CLN2 disease mice

**DOI:** 10.1101/2024.03.29.587276

**Authors:** Keigo Takahashi, Nicholas R. Rensing, Elizabeth M. Eultgen, Sophie H. Wang, Hemanth R. Nelvagal, Steven Q. Le, Marie S. Roberts, Balraj Doray, Edward B. Han, Patricia I. Dickson, Michael Wong, Mark S. Sands, Jonathan D. Cooper

## Abstract

GABAergic interneuron deficits have been implicated in the epileptogenesis of multiple neurological diseases. While epileptic seizures are a key clinical hallmark of CLN2 disease, a childhood-onset neurodegenerative lysosomal storage disorder caused by a deficiency of tripeptidyl peptidase 1 (TPP1), the etiology of these seizures remains elusive. Given that *Cln2^R207X/R207X^* mice display fatal spontaneous seizures and an early loss of several cortical interneuron populations, we hypothesized that those two events might be causally related. To address this hypothesis, we first generated an inducible transgenic mouse expressing lysosomal membrane-tethered TPP1 (TPP1LAMP1) on the *Cln2^R207X/R207X^* genetic background to study the cell-autonomous effects of cell-type-specific TPP1 deficiency. We crossed the TPP1LAMP1 mice with *Vgat-Cre* mice to introduce interneuron-specific TPP1 deficiency. *Vgat-Cre*; TPP1LAMP1 mice displayed storage material accumulation in several interneuron populations both in cortex and striatum, and increased susceptibility to die after PTZ-induced seizures. Secondly, to test the role of GABAergic interneuron activity in seizure progression, we selectively activated these cells in *Cln2^R207X/R207X^* mice using Designer Receptor Exclusively Activated by Designer Drugs (DREADDs) in in *Vgat-Cre*: *Cln2^R207X/R207X^* mice. EEG monitoring revealed that DREADD-mediated activation of interneurons via chronic deschloroclozapine administration accelerated the onset of spontaneous seizures and seizure-associated death in *Vgat-Cre*: *Cln2^R207X/R207X^* mice, suggesting that modulating interneuron activity can exert influence over epileptiform abnormalities in CLN2 disease. Taken together, these results provide new mechanistic insights into the underlying etiology of seizures and premature death that characterize CLN2 disease.

## Introduction

GABAergic interneuron deficits lead to insufficient inhibition or disinhibition, causing an imbalance between excitatory and inhibitory influences within local neural circuits (1–3). Such excitatory/inhibitory imbalance has been implicated in various neurological conditions including epileptic seizures, and several neurodegenerative and psychiatric disorders (4–6). While selective loss or dysfunction of interneurons has been documented in numerous neurological diseases (7–12), such evidence primarily stems from descriptive studies, and direct evidence delineating a causal relationship between interneuron deficits and neurological symptoms remains limited. Filling these knowledge gaps is critical not only for elucidating the functional contribution of inhibitory networks to disease pathogenesis but also for developing potential interneuron-targeted therapeutic strategies.

The neuronal ceroid lipofuscinoses (NCLs) are a group of neurodegenerative lysosomal storage disorders (LSDs) affecting children and young adults (13). CLN2 disease, one of the most common forms of NCLs, is caused by genetic defects in the lysosomal serine protease tripeptidyl peptidase 1 (TPP1), which is encoded by the *CLN2/TPP1* gene (14, 15). Clinically, children affected by CLN2 disease manifest new-onset epileptic seizures between 2 and 4 years of age, followed by ataxia, motor decline, visual loss, and premature mortality (16). Although cerliponase alfa, an FDA-approved disease-modifying enzyme replacement therapy (ERT), now exists (17), this treatment only slows but fails to halt disease progression (18, 19). Notably, managing seizures poses a significant challenge in CLN2 disease clinical care, as seizures are often polymorphic (e.g., generalized tonic-clonic, myoclonic, atonic), progressively worsen and become therapy-resistant as the disease advances (20). Despite the urgent need for more effective treatments, little is known about the cellular mechanisms through which TPP1 deficiency triggers epilepsy and neurodegeneration (21, 22), hindering the identification of novel therapeutic targets and advancement of alternative treatment strategies for CLN2 disease.

Our previous investigation in *Cln2^R207X/R207X^* knock-in mice harboring a common disease-causing mutation (23) revealed a clinically relevant seizure phenotype as a major contributor to premature death (24). Additionally, we also revealed that *Cln2^R207X/R207X^* mice exhibit early loss of several cortical interneuron populations including those positive for parvalbumin (PV), somatostatin (SST), calretinin (CR) and calbindin (CB) (24). While interneuron loss has also been identified in CLN2 patients (25) and a *CLN2^R208X/R208X^* porcine model (26), a key finding from our longitudinal characterization of *Cln2^R207X/R207X^* mice is that this interneuron loss precedes an onset of cortical pyramidal neuron loss and spontaneous seizures (24). These findings prompted us to explore a possible causal link between fatal seizures and cortical interneuron loss in *Cln2^R207X/R207X^* mice.

Studying cell-autonomous effects of TPP1 deficiency is complicated due to TPP1 being secreted from cells and taken up by neighboring cells via the mannose-6-phosphate receptor (M6PR)-mediated endocytosis pathway (27, 28). This process is known as “cross-correction,” and prevents the generation of a cell-autonomous response following cell-type-specific TPP1 deletion. To address this, we generated transgenic mice that express a lysosomal membrane-tethered version of TPP1 (TPP1LAMP1 mice) to suppress cross-correction. TPP1LAMP1 mice are on the *Cln2^R207X/R207X^* genetic background, and ubiquitous expression of the TPP1LAMP1 transgene is designed to suppress their CLN2-disease associated phenotype. Because this transgene is flanked by *LoxP* sites, the Cre-Lox system can be used to produce TPP1 deficiency by excising this transgene in a specific cell type. In this study, crossing TPP1LAMP1 mice with pan-interneuron *Cre* driver mice achieved interneuron-specific TPP1 deficiency. These mice exhibited a shorter lifespan following chemically induced seizures, indicating impaired suppression of fatal seizures. Additionally, we employed a chemogenetic strategy using DREADDs (designer receptors exclusively activated by designer drugs) to selectively activate interneuron activity in *Cln2^R207X/R207X^* mice. Our results demonstrate that chronic DREADD activation of interneurons accelerates the course of spontaneous seizures and associated death in *Cln2^R207X/R207X^* mice. These findings support our hypothesis that interneurons play a mechanistic role in certain aspects of fatal seizure phenotypes in *Cln2^R207X/R207X^* mice.

## Results

### A membrane-tethered in vivo model of TPP1 enables studying the cell-autonomous effects of TPP1 deficiency

Investigating the contribution of interneuron deficits to CLN2 disease pathogenesis required the generation of a new conditional *Tpp1* mutant mouse. To prevent cross-correction of neighboring TPP1-deficient cell types by secreted enzyme, we adopted a membrane-tethering approach that was previously validated in a mouse model of Krabbe disease, which is caused by a deficiency of a different lysosomal enzyme, galactocerebrosidase (GALC) (29). We designed a chimeric fusion protein in which human TPP1 is tethered to the lysosomal membrane by linking TPP1 to the transmembrane domain and cytosolic tail of the lysosomal associated membrane protein 1 (LAMP1) via a 6-glycine linker (Figure 1A). Lentivirus-driven over-expression of the chimeric TPP1LAMP1 in mouse embryonic fibroblasts (MEFs) derived from a Cln2 null mutant mouse (*Cln2*^−/−^) (30) rescued intracellular TPP1 enzymatic activity but did not result in extracellular secretion of TPP1 into the conditioned media or cross-correction in co-cultured *Cln2*^−/−^ MEFs (Figure S1A and B).

**Figure 1.**
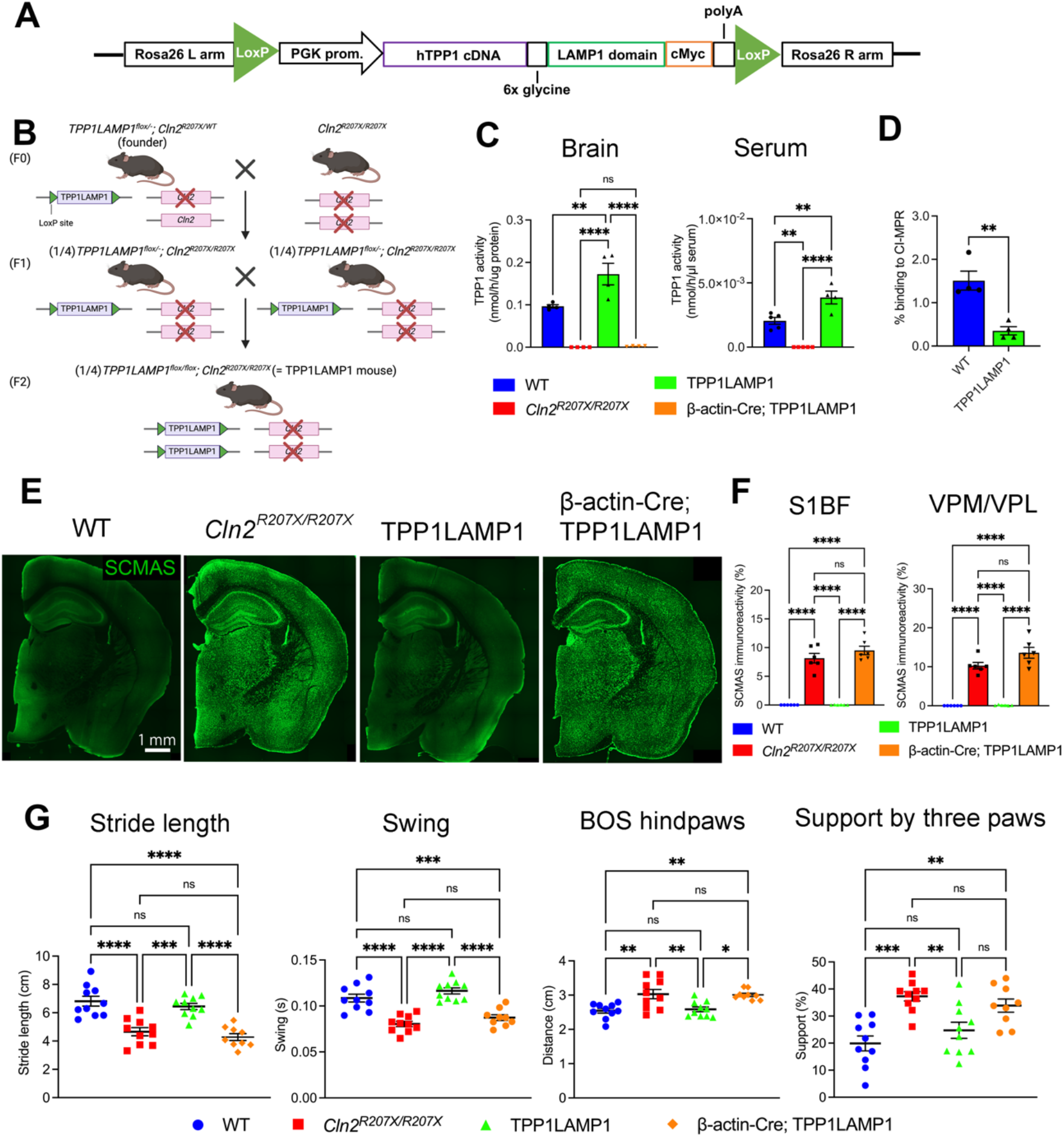
Generation of TPP1LAMP1 mice. (*A*) TPP1LAMP1 construct was created by linking human TPP1 cDNA with the transmembrane domain of LAMP1 via a six-glycine linker under the PGK promoter. The entire construct was flanked by *LoxP* loci. (*B*) Schematic describing the breeding strategy to generate TPP1LAMP1 mice. Floxed-TPP1LAMP1 was inserted into Rosa26 locus, and the founder mice were crossed with *Cln2^R207X/R207X^* mice to achieve double homozygosity for *TPP1LAMP1^flox^* and *Cln2^R207X^*. These mice are designed so that crossing to a specific Cre-driver line will excise the *TPP1LAMP1^flox^* transgene within the cell type of choice to induce TPP1 deficiency due to the *Cln2^R207X/R207X^* genetic background. Mouse drawing was created with BioRender.com. (*C*) TPP1 activity assays in the brain (left) at 15 weeks of age show supraphysiological TPP1 activity in TPP1LAMP1 mice compared to WT mice, and indistinguishable TPP1 activity in β-actin-Cre; TPP1LAMP1 mice from *Cln2^R207X/R207X^* mice (n = 4 per group). TPP1 activity assays in the serum (right) show supraphysiological TPP1 activity in the serum of TPP1LAMP1 mice compared to WT (n = 5 in WT and *Cln2^R207X/R207X^* mice and n = 4 in TPP1LAMP1 mice). (*D*) Binding to the cation-independent mannose-6-phophate receptor (CI-MPR) is significantly reduced in TPP1LAMP1 compared to endogenous WT TPP1 in the brain (n = 4 per group). (*E*) Immunostaining for SCMAS (green) shows the widespread SCMAS accumulation in *Cln2^R207X/R207X^* mice and β-actin-Cre; TPP1LAMP1 mice, but not in WT and TPP1LAMP1 mice at 15 weeks of age. (*F*) Quantitative analysis of SCMAS immunoreactivity via thresholding image analysis in S1BF and VPM/VPL at 15 weeks of age confirms that pathological SCMAS accumulation seen in *Cln2^R207X/R207X^* mice (red bars) is completely rescued in TPP1LAMP1 mice (green bars) and is fully recapitulated in β-actin-Cre; TPP1LAMP1 mice (purple bars) across multiple brain regions (n = 6 per group). (*G*) *CatWalk XT* gait analysis shows significantly shorter stride length and swing duration, wider distance between hind paws (base of support or BOS), and higher proportion of steps supported by 3 feet in 15-week-old *Cln2^R207X/R207X^* mice (red) compared to age-matched WT mice (blue). These gait abnormalities are rescued in TPP1LAMP1 mice (green) and recapitulated in β-actin-Cre; TPP1LAMP1 mice (purple) at the same age (n = 10 in WT, *Cln2^R207X/R207X^*, and TPP1LAMP1 mice and n = 9 in β-actin-Cre; TPP1LAMP1). Dots represent values from individual animals. Values are shown as mean ± SEM. One-way ANOVA with Bonferroni correction in (*C*), (*F*), and (*G*), and unpaired t-test in (*D*). \**P* < 0.05, \*\**P* < 0.01, \*\*\**P* < 0.001, \*\*\*\**P* < 0.0001.

Having validated the function of TPP1LAMP1 protein *in vitro*, we generated a transgenic mouse that ubiquitously expresses *loxP*-flanked TPP1LAMP1 under a phosphoglycerate kinase (PGK) promotor upon the *Cln2^R207X/R207X^* genetic background using CRISPR/Cas9-mediated targeting strategy. The design of this ubiquitous expression of *loxP*-flanked TPP1LAMP1 transgene was intended to normalize disease phenotypes of *Cln2^R207X/R207X^* mice and enable conditional elimination of TPP1 enzymatic activity using the *Cre-loxP* system. After confirming the germline insertion of TPP1LAMP1 transgene into the Rosa26 locus in founder mice (Figure S2A), we crossed the founder mice with *Cln2^R207X/R207X^* to achieve double homozygosity for *TPP1LAMP1^flox^* and *Cln2^R207X^* (*TPP1LAMP1^flox/flox^; Cln2^R207X/R207X^* mice, henceforth referred to as TPP1LAMP1 mice) (Figure 1B). PCR analysis on mRNA-derived cDNA from the liver tissue of TPP1LAMP1 mice confirmed that TPP1LAMP1 is transcribed as a single unit (Figure S2B). TPP1 activity assays in total brain protein extracts also confirmed supraphysiological TPP1 enzymatic activity in TPP1LAMP1 mice *in vivo* compared to WT mice (Figure 1C). However, TPP1 assays in the serum unexpectedly revealed detectable extracellular TPP1 enzymatic activity both in TPP1LAMP1 and WT mice (Figure 1C), in contrast with our aforementioned *in vitro* findings (Figure S1A and B). Because this unanticipated presence of circulating TPP1 could undermine the usefulness of the TPP1LAMP1 mouse as a tool for testing cellular autonomy, we investigated whether these enzymes could bind to cation-independent mannose-6-phosphate receptor (CI-MPR) on the surface of neighboring cells and be delivered into lysosomes. Therefore, we performed an CI-MPR affinity assay on the total brain protein extracts from TPP1LAMP1 and WT mice. The analysis revealed significantly reduced (∼ 77%) affinity of TPP1LAMP1 for the CI-MPR beads compared to endogenous TPP1 (Figure 1D), in line with the previous *in vitro* reports (29). Taken together, these data from *in vivo* samples demonstrate that although membrane tethering does not completely prevent extracellular secretion of TPP1, it significantly restricts cross-correction of the enzyme, validating TPP1LAMP1 mice as an appropriate model to study the cell-autonomous function of TPP1 deficiency.

Next, we verified whether expression of TPP1LAMP1 rescues CLN2 disease-associated phenotypes at 15 weeks (3.5 months) of age, which represents disease end stage for *Cln2^R207X/R207X^* mice. This is an important pre-requisite for demonstrating that the enzyme expressed by our TPP1LAMP1 transgene is biologically active *in vivo*. Storage material accumulation is a pathological hallmark of LSDs (31, 32), and *Cln2^R207X/R207X^* mice display pronounced accumulation of subunit-C of mitochondrial ATPase (SCMAS) throughout the central nervous system (CNS) (24). Immunohistochemistry revealed that SCMAS accumulation is fully prevented in the brain of TPP1LAMP1 mice (Figure 1E), which was further confirmed by quantitative thresholding image analysis within somatosensory pathways including the primary somatosensory barrel cortex (S1BF) and the ventral posterior medial/lateral thalamic nuclei (VPM/VPL), which are the most affected brain regions in *Cln2^R207X/R207X^* mice (Figure 1F). Gait disturbance is also observed in *Cln2^R207X/R207X^* mice at disease end stage (24), and *CatWalk XT* gait analysis showed fully preserved gait performance in TPP1LAMP1 mice (Figure 1G). These results demonstrate that TPP1LAMP1 mice are histologically and behaviorally as healthy as WT mice.

As a final step of validation, we tested whether *Cre-loxP* recombination efficiently excises *loxP*-flanked TPP1LAMP1 transgene, by crossing TPP1LAMP1 mice to *β-actin-Cre* mice, which ubiquitously express *Cre* (*β-actin-Cre^+/-^; TPP1LAMP1^flox/-^; Cln2^R207X/R207X^* mice, henceforth referred to as β-actin-Cre; TPP1LAMP1 mice). PCR analysis targeting the entire TPP1LAMP1 insert at the Rosa26 locus revealed the emergence of a new amplicon that is consistent with the length of the floxed allele (Figure S2C), indicating that the *loxP*-flanked TPP1LAMP1 allele was properly excised. In agreement with this result, TPP1 activity assay in total brain protein extracts revealed dramatically reduced enzymatic activity (2.8% of WT) in β-actin-Cre; TPP1LAMP1 mice, which is comparable to TPP1 activity levels in *Cln2^R207X/R207X^* mice (Figure 1C). Histological analysis and *CatWalk XT* gait analysis showed widespread SCMAS accumulation and gait abnormalities, respectively, in β-actin-Cre; TPP1LAMP1 mice to a similar extent as in *Cln2^R207X/R207X^* mice (Figure 1E, F, and G).

Collectively, these results confirm that ubiquitous *Cre-loxP*-mediated TPP1LAMP1 excision efficiently recapitulates disease phenotypes of *Cln2^R207X/R207X^* mice, validating TPP1LAMP1 mice as an ideal platform to study cell-autonomous effect of conditional TPP1 deficiency *in vivo*.

### Interneuron-specific TPP1 deficiency partially recapitulates neuropathology of CLN2 disease

To elicit TPP1 deficiency in multiple GABAergic interneuron populations, we selected *Vgat-Cre* mice in which *Cre* was inserted downstream of the solute carrier family 32 (GABA vesicular transporter), member 1 gene (*Slc32a1* or *Vgat*) (33). To validate that *Cre* expression was present within these interneuron populations in *Vgat-Cre* mice, we first crossed *Vgat-Cre* mice to *Ai14* mice, a reporter strain that expresses tdTomato in the presence of *Cre* (34). Histological analysis of the brain sections from *Vgat-Cre^+/-^; Ai14^+/-^*mice revealed that major inhibitory neuron populations including PV-, SST-, and CR-positive interneurons within S1BF co-localize with tdTomato (Figure S3A and B). In the S1BF, 32.9%, 26.1%, and 8.3% of tdTomato-positive cells are PV-, SST-, and CR-positive neurons, respectively (Figure S3C), and this composition is largely consistent with the previous literature (35, 36). We also found that COUP-TF-interacting protein 2 (CTIP2)-positive interneurons within caudate putamen (CPu) in the striatum, known as medium-sized spiny neurons (MSNs), also co-localize with the tdTomato signal (Figure S3A and B). Taken together, these results confirm that *Vgat-Cre* effectively targets interneurons in multiple brain regions.

Next, we crossed TPP1LAMP1 mice to *Vgat-Cre* mice to introduce interneuron-specific TPP1 deficiency (*Vgat-Cre^+/-^; TPP1LAMP1^flox/flox^; Cln2^R207X/R207X^* mice, henceforth referred to as Vgat-Cre; TPP1LAMP1 mice). Immunostaining for SCMAS, the key pathological hallmark of CLN2 disease, in the brain of Vgat-Cre; TPP1LAMP1 mice at 15 weeks of age showed scattered distribution of SCMAS-positive cells across the brain (Figure 2A and B). To confirm whether these SCMAS-positive cells are interneurons, we counter-stained brain sections of Vgat-Cre; TPP1LAMP1 mice for CTIP2, PV, and SST. Co-immunostaining revealed that SCMAS accumulates in a subset of CTIP2-positive MSNs within CPu and PV- and SST-positive interneurons within S1BF (Figure 2C – E), suggesting successful TPP1 deletion in these interneuron populations. To investigate potential loss of those interneuron populations in Vgat-Cre; TPP1LAMP1 mice, we performed unbiased stereological counting on immunostained brain sections. Statistical analysis revealed a significant loss of CTIP2-positive MSNs within CPu in Vgat-Cre;TPP1LAMP1 mice compared to age-matched TPP1LAMP1 mice, which is indistinguishable from age-matched *Cln2^R207X/R207X^* mice (Figure 2F). However, the same unbiased stereological analysis within S1BF reveals no significant reduction of PV- or SST-positive neurons in Vgat-Cre; TPP1LAMP1 mice compared to age-matched TPP1LAMP1 mice (Figure 2F), suggesting a subpopulation-dependent impact of TPP1 deficiency on interneuron loss in Vgat-Cre; TPP1LAMP1 mice.

**Figure 2.**
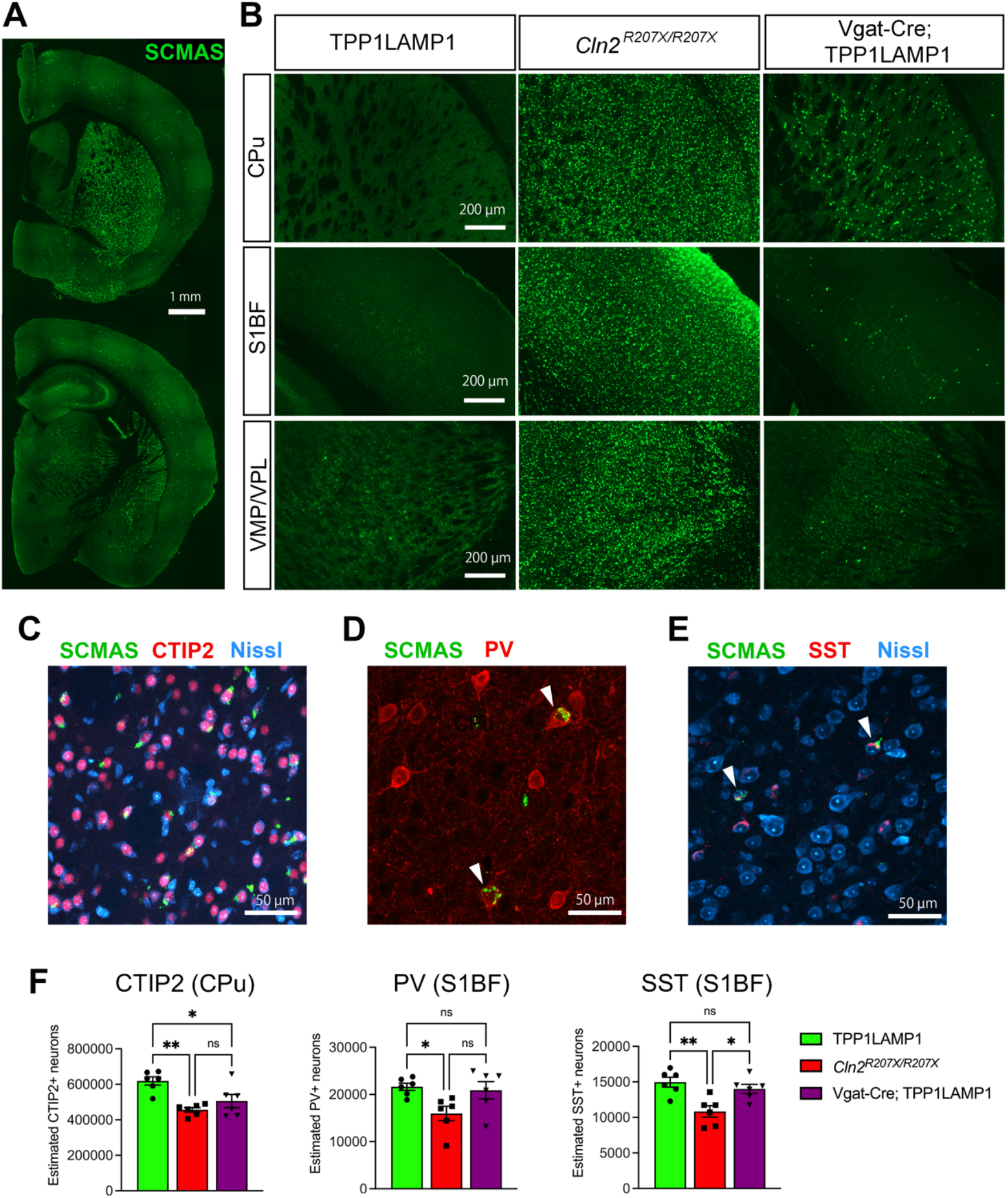
Interneuron-specific TPP1 deficiency leads to storage material accumulation and interneuron loss. (*A*) Immunostaining for SCMAS (green) in the brain of Vgat-Cre; TPP1LAMP1 at 15 weeks of age shows scattered distribution of SCMAS-positive cells across the brain. (*B*) Comparison of immunostained brain sections for SCMAS (green) between TPP1LAMP1, *Cln2^R207X/R207X^*, and Vgat-Cre; TPP1LAMP1 mice across multiple brain regions reveals partial storage material accumulation in Vgat-Cre; TPP1LAMP1 compared to *Cln2^R207X/R207X^* mice with the greatest extent within caudate putamen (CPu) at 15 weeks of age. (*C*) Co-immunostaining for SCMAS (green), CTIP2 (red), and Nissl (cyan) reveals storage material accumulation in a subset of CTIP2-positive medium spiny neurons within CPu in Vgat-Cre; TPP1LAMP1 mice. (*D*) Co-immunostaining for SCMAS (green) and PV (red) reveals storage material accumulation in a subset of PV-positive interneurons within S1BF cortex in Vgat-Cre; TPP1LAMP1 mice. (*E*) Co-immunostaining for SCMAS (green), SST (red), and Nissl (cyan) reveals storage material accumulation in a subset of SST-positive interneurons within S1BF cortex in Vgat-Cre; TPP1LAMP1 mice. (*F*) Unbiased stereological counts of immunostained neuron populations within CPu reveals a significant loss of CTIP2-positive medium spiny neurons in Vgat-Cre: TPP1LAMP1 mice to the comparable extent to that in *Cln2^R207X/R207X^* mice at 15 weeks of age. The same unbiased stereological analysis within S1BF cortex reveals no significant loss of PV or SST-positive neurons in Vgat-Cre; TPP1LAMP1 mice compared to those in TPP1LAMP1 mice at 15 weeks of age. Dots represent values from individual animals. Values are shown as mean ± SEM (n = 6 mice per group). One-way ANOVA with Bonferroni correction. \**P* < 0.05, \*\**P* < 0.01.

Localized glial activation of both astrocytes and microglia is another pathological hallmark of NCLs, including CLN2 disease (37). To assess whether interneuron-specific TPP1 deficiency triggers glial activation, we performed immunohistochemistry on brain sections of 15-week-old Vgat-Cre; TPP1LAMP1 mice for glial fibrillary acidic protein (GFAP), a marker for astrogliosis, and CD68, a marker for microglial activation (37, 38). Immunohistological analysis revealed a marked increase in both GFAP and CD68 immunoreactivities in age-matched *Cln2^R207X/R207X^* mice across CPu, S1BF, and VPM/VPL, but no significant increase was observed within any of these brain regions of Vgat-Cre; TPP1LAMP1 mice compared to age-matched TPP1LAMP1 mice (Figure 3A and B). These results suggest that TPP1-deficient interneurons do not trigger a non-cell-autonomous neuroimmune response associated with CLN2 disease.

**Figure 3.**
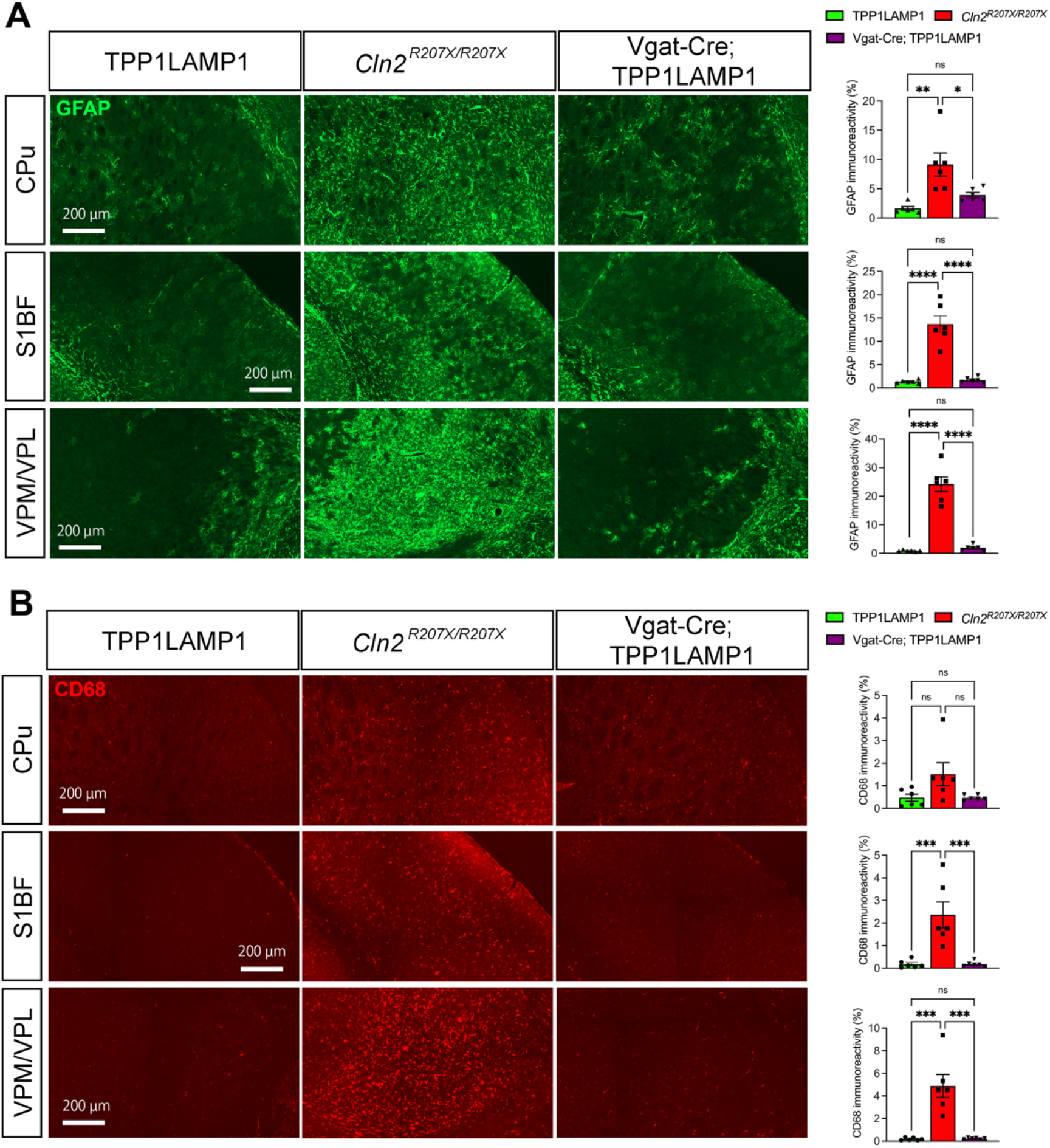
Interneuron-specific TPP1 deficiency does not trigger the neuroimmune response associated with CLN2 disease. Immunostaining for GFAP (*A*, green) and CD68 (*B,* red) and quantitative analysis of their immunoreactivity via thresholding image analysis in CPu, S1BF, and VPM/VPL at 15 weeks of age shows the marked increase in the intensity of GFAP and CD68 immunoreactivities in *Cln2^R207X/R207X^* mice (red bars), but no significant increase in Vgat-Cre; TPP1LAMP1 mice (purple bars) compared to age-matched TPP1LAMP1 mice (green bars). Dots represent values from individual animals. Values are shown as mean ± SEM (n = 6 mice per group). One-way ANOVA with Bonferroni correction. \**P* < 0.05, \*\**P* < 0.01. \*\*\**P* < 0.001.

### Interneuron-specific TPP1 deficiency increases susceptibility to sudden death secondary to PTZ-induced seizures

We then investigated whether Vgat-Cre; TPP1LAMP1 mice demonstrate any neurological phenotypes including the gait disturbance and seizure phenotypes associated with CLN2 disease. *CatWalk XT* gait analysis revealed no significant deterioration of gait parameters in 15-week-old Vgat-Cre; TPP1LAMP1 mice compared to age-matched TPP1LAMP1 mice (Figure 4A). The same gait analysis was repeated at an older age (25 weeks old), yet no significant change was observed between Vgat-Cre; TPP1LAMP1 and TPP1LAMP1 mice (data not shown). These data suggest no apparent impact of interneuron-specific TPP1 deficiency on gait performance.

**Figure 4.**
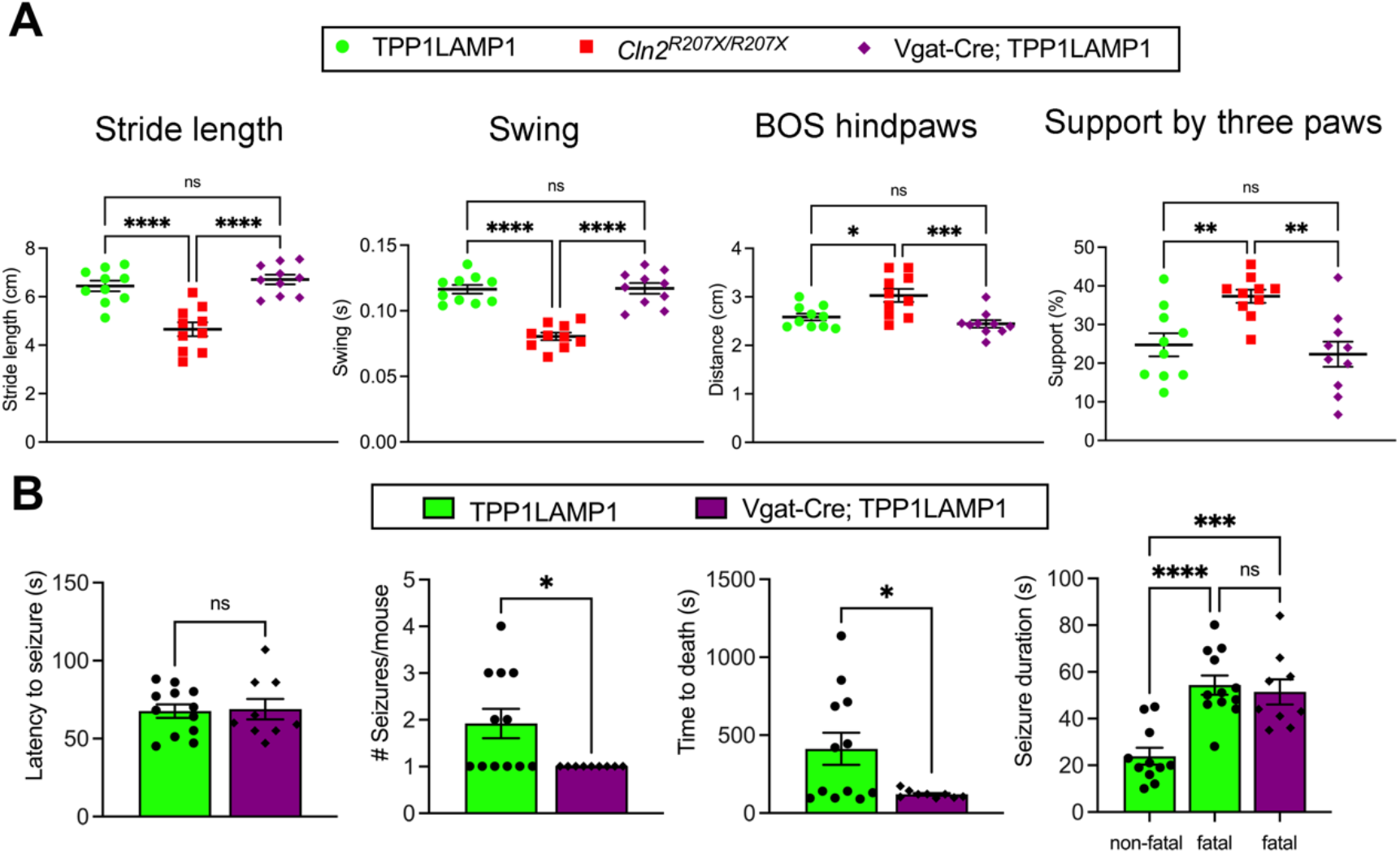
Interneuron-specific TPP1 deficiency increases susceptibility to sudden death secondary to PTZ-induced seizures. (*A*) *CatWalk XT* gait analysis reveals preserved gait performance of Vgat-Cre; TPP1LAMP1 mice (purple) compared to TPP1LAMP1 and *Cln2^R207X/R207X^* mice at 15 weeks of age (n = 10 per group). (*B*) Seizures were induced by intraperitoneal injection of PTZ 75mg/kg in TPP1LAMP1 (n = 12, green bars) and Vgat-Cre; TPP1LAMP1 (n = 9, purple bars) mice at 25 weeks of age. There is no significant deference in the latency to tonic-clonic seizures between two groups. Vgat-Cre; TPP1LAMP1 mice exhibit a significantly lower number of tonic-clonic seizures per mouse and a significantly shorter time to death compared to TPP1LAMP1 mice. The average duration of non-fatal tonic-clonic seizures in TPP1LAMP1 mice is significantly shorter than the average duration of fatal tonic-clonic seizures in TPP1LAMP1 mice and Vgat-Cre; TPP1LAMP1 mice. Dots represents values from individual animals. Values are shown as mean ± SEM. One-way ANOVA with Bonferroni correction in (A) and unpaired t-test in (*B*). \**P* < 0.05, \*\**P* < 0.01. \*\*\**P* < 0.001, \*\*\*\**P* < 0.0001.

Next, we performed long-term electroencephalogram (EEG) monitoring on Vgat-Cre; TPP1LAMP1 mice. We previously reported that *Cln2^R207X/R207X^* mice start to display abnormal background EEG activity such as epileptiform spikes and spike bursts from approximately 12 weeks of age onwards and subsequently develop spontaneous tonic-clonic seizures with a median onset of 15 weeks of age (24). However, EEG monitoring in Vgat-Cre; TPP1LAMP1 mice revealed no apparent interictal epileptiform background abnormalities or spontaneous seizures up to 30 weeks of age. Therefore, we induced seizures in both TPP1LAMP1 (n=12) and Vgat-Cre; TPP1LAMP1 mice (n=9) at 25 weeks of age by intraperitoneal (IP) injection of pentylenetetrazole (PTZ) at 75mg/kg of body weight. This approach provided a more sensitive measure to interrogate whether interneuron-specific TPP1 deficiency affects seizure susceptibility. Our analysis revealed no significant difference in the average latency to tonic-clonic seizures (Figure 4B), suggesting that the threshold for seizure initiation is not altered in Vgat-Cre; TPP1LAMP1 mice. However, while half (6/12) of the TPP1LAMP1 mice died after a single seizure and the remaining mice died after experiencing multiple seizures, in contrast all (9/9) of the Vgat-Cre; TPP1LAMP1 mice died immediately after a single tonic-clonic seizure (p = 0.0186, Fisher’s exact test). This resulted in a significantly lower average number of tonic-clonic seizures before dying and a significantly shorter average time of death in Vgat-Cre; TPP1LAMP1 mice compared to TPP1LAMP1 mice (Figure 4B). Furthermore, we found that the average duration of non-fatal tonic-clonic seizures in TPP1LAMP1 mice is significantly lower than the average duration of fatal tonic-clonic seizures in TPP1LAMP1 mice and Vgat-Cre; TPP1LAMP1 mice. Taken together, these results suggest that Vgat-Cre; TPP1LAMP1 mice are less capable of terminating PTZ-induced seizures, and thus more prone to die following PTZ-induced seizures compared to TPP1LAMP1 mice.

### Chemogenetic activation of interneurons exacerbates seizure phenotypes in Cln2^R207X/R207X^ mice

As a complementary approach to investigate the contribution of interneurons to seizure phenotypes in *Cln2^R207X/R207X^* mice, we chemogenetically activated interneurons using hM3Dq, a modified form of the human M3 muscarinic (hM3) receptor classified as one of DREADDs. *Cre*-dependent AAV9-hSyn-DIO-hM3Dq-mCherry or AAV9-hSyn-DIO-mCherry (control, n = 8 per group) was intracerebroventricularly (ICV) injected into neonatal (P1-2) *Vgat-Cre^+/-^; Cln2^R207X/R207X^* mice to transduce hM3Dq-DREADD in interneurons across the brain, predominantly in the cortex and striatum (Figure 5A and B). At 78 days (11 weeks 0 days) of age, which is just before *Cln2^R207X/R207X^* mice start to display epileptiform abnormalities, we started to chronically administer 10 µg/ml of deschloroclozapine (DCZ), a recently described DREADD ligand that appears to be more potent and specific compared to other classic DREADD ligands (39, 40), via drinking water in both treatment groups (Figure 5A). While DREADD ligands have been more commonly administered by intraperitoneal injection, this non-invasive oral administration route of DCZ was selected to minimize any additional handling stress that could potentially induce seizures in *Cln2^R207X/R207X^* mice. Long-term EEG monitoring was also initiated at 78 days of age and continued until the mice died. The hM3Dq-DREADD activated group showed a significantly earlier onset of spontaneous seizures, along with earlier-onset of interictal abnormalities including epileptiform spikes and bust-suppression pattern, compared to the mCherry-expressing control group (Figure 5C, D, and F). This led to significantly earlier death in the hM3Dq-DREADD activated group compared to the mCherry control group (Figure 5C and E). Taken together, these results indicate that chemogenetic chronic activation of interneurons exacerbates epileptic seizures and accelerates seizure-related death in *Cln2^R207X/R207X^* mice, providing additional evidence in support of the involvement of interneurons in epileptogenesis associated with CLN2 disease.

**Figure 5.**
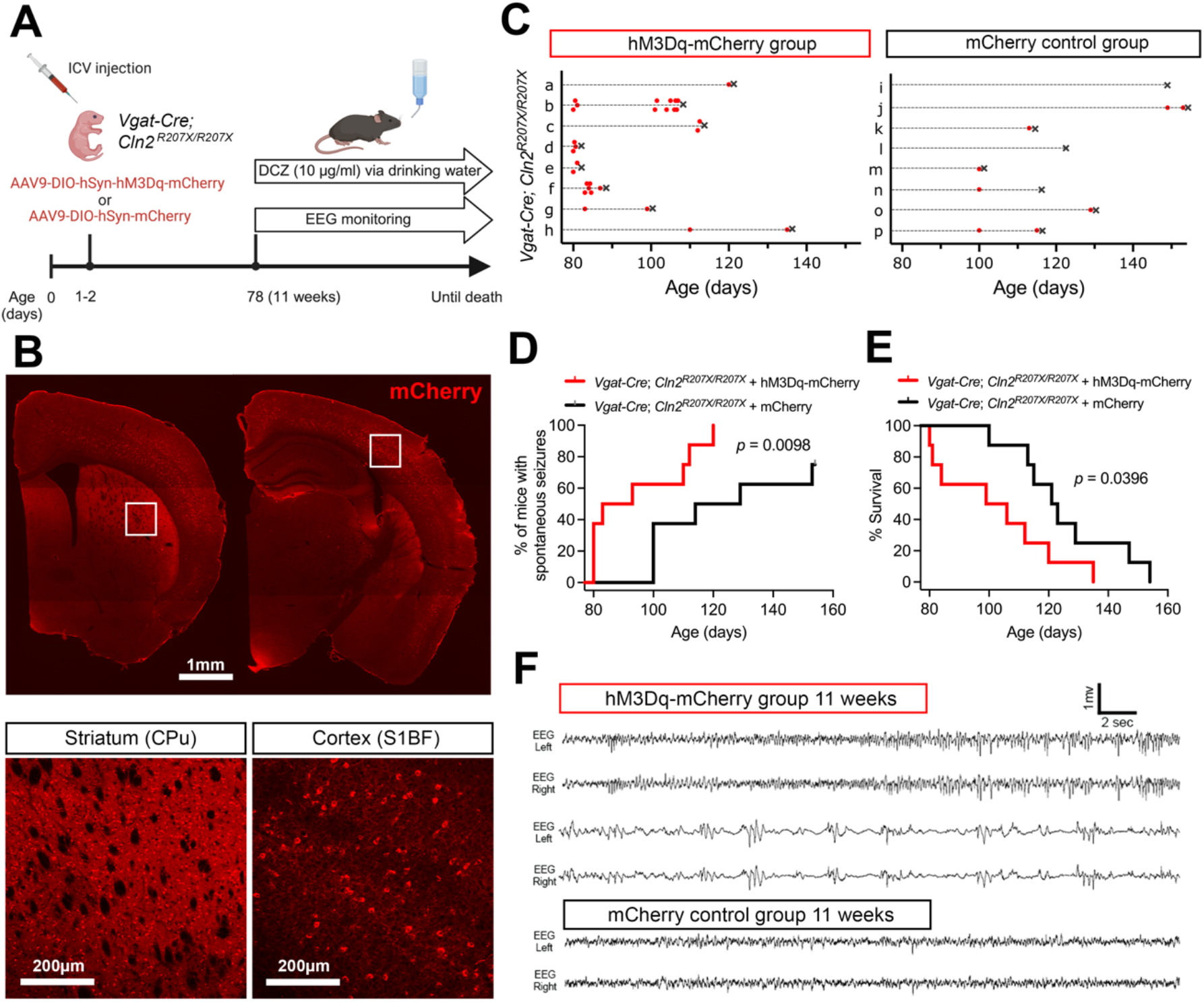
Chemogenetic activation of interneurons exacerbates seizure phenotypes in *Cln2^R207X/R207X^* mice. (*A*) Schematic of experimental design. The image was created with BioRender.com. (*B*) Fluorescence images depict widespread transduction of intracerebroventricularly delivered AAV9-hSyn-DIO-hM3Dq-mCherry (red) in *Vgat-Cre*; *Cln2^R207X/R207X^* mouse brain at 6 weeks of age, predominantly in the cortex and striatum. Lower magnification images (top) and higher magnification images focused on the striatum (CPu) and cortex (S1BF, bottom). (*C*) Time course of seizures (red dots) and deaths (black Xs) in hM3Dq-mCherry-expressing (top) and mCherry-expressing (bottom) *Vgat-Cre*; *Cln2^R207X/R207X^* mice. EEG recording reveals a significantly earlier onset of spontaneous seizures (*D*) and premature death (*E*) in hM3Dq-mCherry-expressing *Vgat-Cre*; *Cln2^R207X/R207X^* mice upon chronic DCZ administration compared to mCherry-expressing control mice (n = 8 per group). Log-rank (Mantel-Cox) test. (*F*) Representative EEG traces show an increased frequency of abnormal spikes (top) and burst-suppression activity (middle) in hM3Dq-mCherry-expressing *Vgat-Cre*; *Cln2^R207X/R207X^* mice during the first week of DCZ administration compared to mCherry-expressing *Vgat-Cre*; *Cln2^R207X/R207X^* mice (bottom).

### Chemogenetic activation of interneurons alters microglial activation and astrocytic GABA levels in Cln2^R207X/R207X^ mice

Observing how chemogenetic activation of interneurons alters seizure phenotypes in *Cln2^R207X/R207X^* mice raised the question of whether interneuron activity also affects the disease-associated neuropathological phenotypes of *Cln2^R207X/R207X^* mice. Thus, we extended our analysis to investigate the neuroimmune responses at a histological level in DREADD-treated *Vgat-Cre^+/-^; Cln2^R207X/R207X^* mice. For this purpose, we treated *Vgat-Cre^+/-^; Cln2^R207X/R207X^* mice with DCZ for 72 hours at 11 weeks of age (Figure 6A). To achieve a tighter control of daily DCZ consumption, we utilized the micropipette-guided administration method (41) rather than administration via drinking water to precisely administer 500µg/kg/day of DCZ in condensed milk to *Vgat-Cre^+/-^; Cln2^R207X/R207X^* mice (Figure 6A). Immunohistological analysis revealed a significant reduction of CD68 immunoreactivity in the hM3Dq-DREADD activated group across CPu, S1BF, and VPM/VPL compared to mCherry control group, whereas no significant change in GFAP immunoreactivity was observed between two groups (Figure 6B).

**Figure 6.**
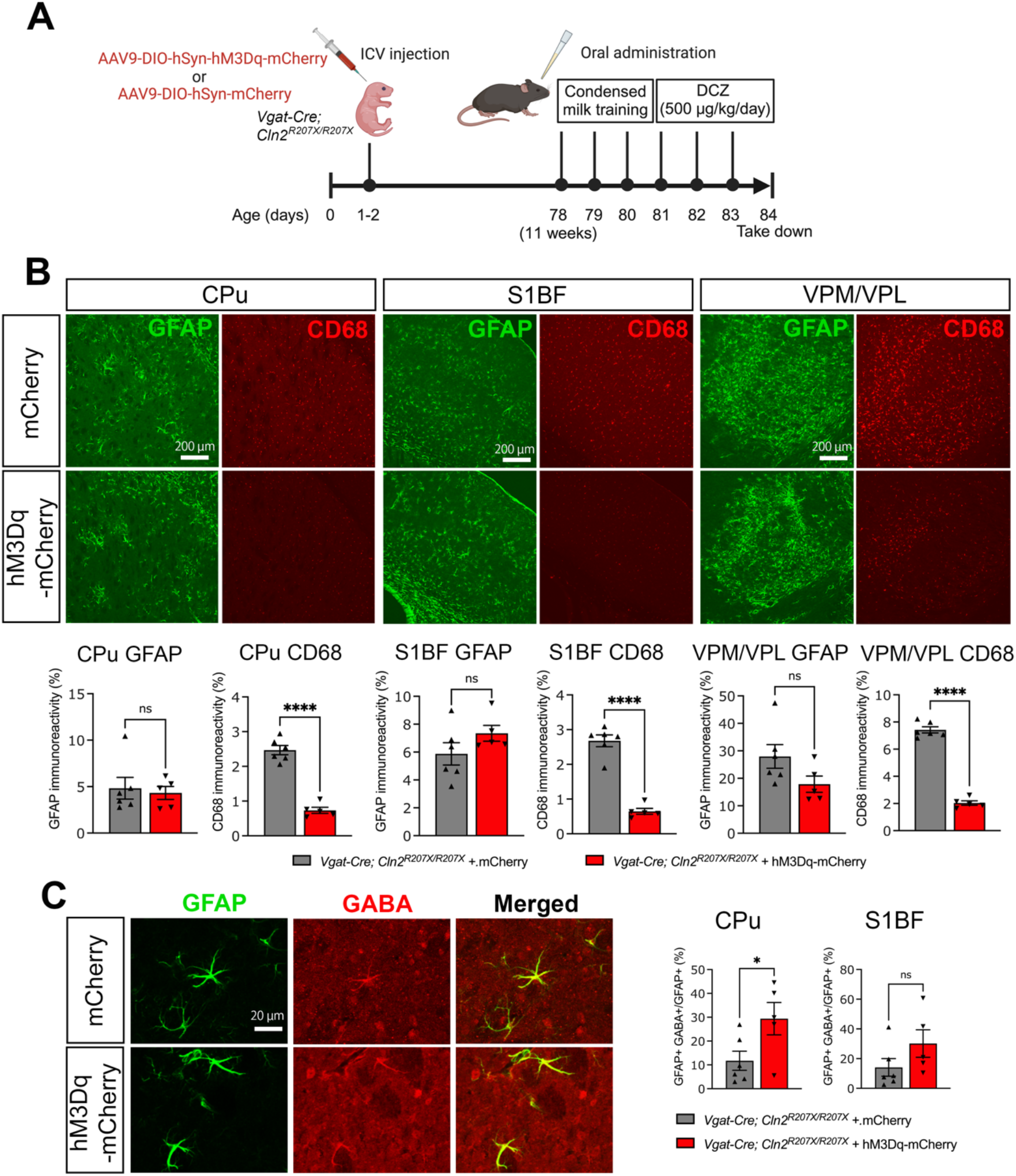
Chemogenetic activation of interneurons alters microglial activation and astrocytic GABA levels in *Cln2^R207X/R207X^* mice. (*A*) Schematic of experimental design. The image was created with BioRender.com. (*B*) Immunostaining for GFAP (green) and CD68 (red) and quantitative analysis of their immunoreactivity via thresholding image analysis reveals a significantly reduced CD68 immunoreactivity in hM3Dq-mCherry-expressing *Vgat-Cre*; *Cln2^R207X/R207X^* mice (n = 5, red bars) upon DCZ administration across CPu, S1BF, and VPM/VPL with no change in GFAP immunoreactivity compared to mCherry-expressing control mice (n = 6, gray bars). (*C*) Immunostaining for GFAP (green) and GABA (red) and co-localization analysis on the confocal images reveals a significantly increased GABA immunoreactivity in GFAP-positive astrocytes within CPu of hM3Dq-mCherry-expressing *Vgat-Cre*; *Cln2^R207X/R207X^* mice (n = 5, red bars) upon DCZ administration compared to mCherry-expressing control mice (n = 6, gray bars). There is a similar increase in the GABA immunoreactivity in GFAP-positive astrocytes within S1BF, but it is not statistically significant. Dots represents values from individual animals. Values are shown as mean ± SEM. Unpaired t-test. \**P* < 0.05, \*\**P* < 0.01. \*\*\**P* < 0.001, \*\*\*\**P* < 0.0001.

While chemogenetic manipulation of interneuron activity appeared to have no effect upon astrogliosis, considering the recent evidence suggesting astrocytic regulation of GABA (42, 43), we next explored whether astrocytic GABA expression is altered in the brain of *Cln2^R207X/R207X^* mice and whether that can be affected by chemogenetic manipulation of interneuron activity. Co-immunostaining for GFAP and GABA revealed pronounced co-localization of these two signals within CPu, S1BF, and VPM/VPL as early as 12 weeks (3 months) of age (Figure S4), suggesting increased levels of GABA in GFAP-positive reactive astrocytes. We then performed GFAP/GABA co-immunostaining on brain sections of hM3Dq-mCherry or mCherry-expressing *Vgat-Cre^+/-^; Cln2^R207X/R207X^* mice treated with DCZ for 72 hours, focusing upon CPu and S1BF as regions where AAV9-mediated hM3Dq transduction primarily occurred. Co-localization analysis on confocal images revealed a significant increase in the co-localization of GFAP and GABA within GFAP-positive structures in the CPu, and a trend toward an increase within S1BF although it was not statistically significant (Figure 6C). Taken together, these data indicate that chemogenetic enhancement of interneuron activity increases local astrocytic GABA levels in *Cln2^R207X/R207X^* mice, suggesting a novel interaction between interneurons and astrocytes in the context of CLN2 disease.

## Discussion

In this study, we have generated and demonstrated the utility of TPP1LAMP1 mice in which cross-correction is prevented as a valuable platform for investigating the cell-autonomous effects of conditional TPP1 deficiency. Cross-correction of lysosomal enzymes between neighboring cells involves both secretion into the extracellular space and subsequent endocytosis via the M6PR pathway (44). Based on our *in vitro* data we initially assumed that membrane tethering would prevent TPP1 from being extracellularly secreted, and the unexpected detection of TPP1 enzymatic activity in the serum of TPP1LAMP1 mice prompts speculation regarding its mechanism. Since a significant pool of LAMP1 traffics via the plasma membrane en route to the lysosome (45), it is very likely that TPP1LAMP1 follows the same trafficking route in cells. Considering that TPP1 has both exopeptidase and endopeptidase functions (46–49), one possibility is that the TPP1LAMP1 which is normally confined to the lysosomal lumen undergoes some autocatalytic cleavage either inside the lysosome or at the cell surface, releasing free TPP1 into the extracellular space. Alternatively, considering that 5 – 20% of newly synthesized lysosomal enzymes escape binding to M6PR and become secreted in the non-tumor cells (50), it is plausible to speculate that TPP1LAMP1 might be prone to extracellular secretion due to altered M6P modification. Another possibility is that TPP1LAMP1 might exist inside exosomes or extracellular vesicles (EVs) released in the extracellular space and evidence suggests that LAMP1 is present in neural-derived plasma exosomes (51). Nevertheless, both TPP1LAMP1 and previously studied GALCLAMP1 (29) exhibit significantly reduced affinity to CI-M6PR compared to WT native enzymes. This suggests that membrane tethering of these lysosomal enzymes markedly alters their mannose-6-phophorylation, thus evading recognition by, and binding to, the cell surface-expressed M6PR, thereby preventing their usual ability to cross correct neighboring cells. Our data demonstrating the accumulation of SCMAS, a sensitive marker for TPP1 deficiency in neurons, within interneurons of Vgat-Cre; TPP1LAMP1 mice, despite being surrounded by other phenotypically healthy cell types, provides further support that TPP1 cross-correction is effectively prevented in TPP1LAMP1 mice. Therefore, despite the unexpected presence of extracellular TPP1 activity in serum, TPP1LAMP1 mice remain a valid model for assessing conditional TPP1 deficiency.

We included CPu, the largest area of murine striatum, in our histological analysis due to observed pronounced SCMAS accumulation in many striatal cells in Vgat-Cre; TPP1LAMP1 mice. GABAergic MSNs account for approximately 95% of striatal neurons in rodents (52), receiving glutamatergic inputs from the cortex and thalamus, as well as dopaminergic inputs from the substantia nigra pars compacta (53). Loss or dysfunction of these striatal MSNs has been implicated in movement disorders such as Parkinson’s disease and Huntington’s disease (54, 55). Although neuron loss and glial activation in the striatum had already been documented in autopsy brain tissues from CLN2 patients (25), striatal pathology has been largely overlooked in CLN2 disease research using animal models. Our findings present the first evidence of CTIP2-positive MSN loss, along with pronounced astrogliosis and microglial activation in mouse models of CLN2 disease. Interestingly, the cell-autonomous loss of MSNs within CPu of Vgat-Cre; TPP1LAMP1 mice did not cause astrogliosis or microglial activation locally, suggesting that CLN2 disease-associated glial activation might also occur autonomously rather than as a secondary response to neuron loss, at least within this brain region. With TPPLAMP1 mice theoretically enabling us to introduce glial-specific TPP1 deficiency, it will be important to explore potential cell-autonomous effects of TPP1 deficiency in different types of glia and its contribution to other phenotypes in *Cln2^R207X/R207X^* mice. Notably, CLN2 patients often exhibit complex movement disorder phenotypes, including non-epileptic myoclonus, dystonia, spasticity, chorea, athetosis, and tremors (56), suggesting an underappreciated involvement of striatal pathology. A better understanding of movement disorder etiology and potential striatal involvement will be crucial for devising therapeutic approaches targeting a wide range of neurological symptoms of CLN2 disease.

Epileptiform activity can be divided into at least three distinct stages: initiation, propagation, and termination (57). Vgat-Cre; TPP1LAMP1 mice did not develop spontaneous seizures during our study period, indicating that interneuron-specific TPP1 deficiency is by itself not sufficient to initiate seizures. To further explore the potential role of TPP1-deficient interneurons in seizure susceptibility, we induced acute generalized seizures in Vgat-Cre; TPP1LAMP1 mice via a single IP injection of PTZ, a commonly used method for both mechanistic studies and preclinical evaluation of anti-epileptic drugs (58). Our findings revealed no significant difference in latency to seizure onset between Vgat-Cre; TPP1LAMP1 mice and TPP1LAMP1 control mice. Although many TPP1LAMP1 mice experienced multiple generalized seizures of shorter duration before advancing to fatal long-lasting seizures, we found that Vgat-Cre; TPP1LAMP1 mice were significantly more likely to die following a single episode of generalized long-lasting seizures. These results suggest that interneurons may play a critical role in terminating seizures rather than initiating them in CLN2 disease. Importantly, sudden death following seizures is a distinctive feature of *Cln2^R207X/R207X^* mice (24) compared to *Cln1^−/−^* mice, modeling CLN1 disease, which develop similar spontaneous seizures without experiencing seizure-related sudden death (59). Therefore, although the nature of PTZ-induced seizures may differ considerably from the nature of chronic spontaneous seizures of *Cln2^R207X/R207X^* mice, our findings provide novel insights into the mechanistic role of interneurons in the fatal seizure phenotypes of CLN2 disease.

Recent technological advancements in chemogenetics and optogenetics have enabled the study of causality between interneuron activity and seizures in animal models, revealing the protective contribution of interneuron activities to various seizure types (60, 61). Initially, we anticipated that hM3Dq-DREADD activation of interneurons would attenuate spontaneous seizures in *Cln2^R207X/R207X^* mice. However, our results robustly showed an opposite effect; chemogenetic activation of interneurons in *Cln2^R207X/R207X^* mice exacerbated their fatal seizure phenotypes. Another study has reported that optogenetic activation of hippocampal PV- and SST-positive interneurons increases chemically induced epileptic discharges in wild type mice (62). Additionally, silencing PV-positive interneurons in the primary motor cortex has been shown to reduce the duration of optogenetically induced seizures (63). These findings suggest a bimodal role of interneuron activity in seizures, dependent upon interneuron sub-populations, brain regions, seizure types, and disease status. We used *Vgat-Cre* mice to simultaneously target multiple interneuron populations in our experiments. This was because interneuron loss has been confirmed to occur to a similar extent in at least five different sub-populations (PV-, SST-, CR-, and CB-positive cortical interneurons and CTIP2-positive striatal MSNs) in *Cln2^R207X/R207X^* mice (24). While this choice of *Cre* driver line was a logical initial step to establish the involvement of interneurons in CLN2 disease, it will be informative to identify the individual interneuron sub-population(s) that contribute most to seizure phenotypes in *Cln2^R207X/R207X^* mice.

It is important to note that while most previous studies have utilized an acute DREADD approach to study chemically induced seizures, our experiment instead employed a chronic DREADD approach. This approach allowed us to monitor the long-term effects of DREADD activation of interneurons on naturally occurring spontaneous seizures in *Cln2^R207X/R207X^* mice over time. As such, caution should be exercised comparing our findings to those derived from acute DREADD experiments. Nevertheless, chronic DREADD studies are becoming increasingly common due to the recent developments in selective, potent, and long-acting DREADD ligands such as DCZ, as well as non-invasive ligand administration routes (64–66). However, it should be pointed out that there are several potential technical concerns associated with chronic DREADD studies, including tolerance and desensitization to the ligand (60). In particular, chronic DREADD activation of interneurons may cause “ionic plasticity,” a phenomenon where continuous activation of interneurons leads to a massive Cl^-^ influx, resulting in a qualitative change in GABAAR-mediated signaling from hyperpolarization (inhibition) to depolarization (excitation) (67). Such depolarized GABAergic activity has been observed in a study where hM3Dq-DREADD activation of interneurons within subiculum for 3 days exacerbated electrographic seizures during the chronic phase of chemically induced epilepsy in *Vgat-Cre* mice (68), suggesting a pro-convulsant effect of ionic plasticity in chronically activated interneurons. Therefore, a functional conversion of interneurons from inhibition to excitation might explain the seizure exacerbation in *Cln2^R207X/R207X^* mice upon chronic DREADD activation of interneurons, and future electrophysiological studies would provide more mechanistic insights into this possibility.

Although localized activation of both microglia and astrocytes has been widely used as a measure of the efficacy of experimental therapies across many NCLs, its pathological role in neurodegeneration remains unclear (37). Despite the pro-convulsant effect of hM3Dq-DREADD activation of interneurons in *Cln2^R207X/R207X^* mice, 72-hour DREADD activation of interneurons attenuated CD68-positive microglial activation, typically interpreted as improvement in disease phenotype. It is known that microglial processes make contact with GABAergic synapses and are involved in synaptic pruning and modulating neurotransmission (69). Emerging evidence also suggests that microglia express GABABRs and sense inhibitory neuronal activity during early developmental stages (70, 71). While little is known about microglial-interneuron interactions in the young adult brain, our finding suggest that microglial reactivity associated with CLN2 disease might be directly influenced by interneuron activity via GABA.

In contrast, hM3Dq-DREADD activation of interneurons did not alter GFAP-positive astrogliosis, but instead significantly increased the levels of GABA within reactive astrocytes in the striatum. Astrocytes are both GABAergic and GABAceptive, playing an important role in GABA metabolism in the brain (42, 43). Therefore, the significant increase in astrocytic GABA upon hM3Dq-DREADD activation of interneurons suggests enhanced astrocytic uptake of GABA that is increasingly released from activated interneurons. Recent studies have demonstrated abnormal GABA increases in reactive astrocytes in various neurological conditions such as Alzheimer’s disease (72), kainic acid-induced epilepsy (73, 74), and stroke (75). To our knowledge, our results provide the first evidence of astrocytic GABA increase in any LSD model. These increases in astrocytic GABA may explain our previous findings of sustained GABA concentration in total protein extracts from the cortex of 3-month-old *Cln2^R207X/R207X^* mice compared to WT mice, despite the profound loss of interneurons (24). While it remains unclear whether astrocytic GABA increase in CLN2 disease serves as a compensatory mechanism in response to interneuron loss, a better understanding of its pathological impact might offer an alternative therapeutic target.

In summary, we have generated a novel TPP1LAMP1 mouse to investigate the cell-autonomous effects of conditional TPP1 deficiency. The TPP1 enzyme present in these mice has poor affinity for M6PR, rendering it incapable of cross-correction, confirming the utility of these mice as a valuable platform for addressing cell-autonomous effects of TPP1 deficiency. Interestingly, while interneuron-specific TPP1 deficiency did not initiate seizures, it rendered mice more susceptible to death secondary to PTZ-induced seizures, suggesting a critical role of interneurons in seizure termination in CLN2 disease. Furthermore, chronic DREADD activation of interneurons exacerbated seizure phenotypes, underscoring the complexity of interneuron involvement in seizures associated with CLN2 disease. Additionally, our findings indicated a direct influence of interneuron activity on microglial reactivity via GABA, while astrocytic GABA increase in CLN2 disease may represent a compensatory mechanism in response to interneuron loss. Our results highlight the intricate interplay between interneurons, glia, and seizures in CLN2 disease, providing valuable insights for the development of targeted therapeutic interventions.

## Materials and Methods

### Mice

*Cln2^R207X/R207X^* (JAX: 030696) (76), *Vgat-Cre* (JAX: 028862) (33), *β-actin-Cre* (JAX: 033984) (77), Ai14 (JAX: 007914) (34), and WT mice were maintained on a C57Bl/6J background and housed in an animal facility at Washington University School of Medicine (St. Louis, MO) under a 12hr light/dark cycle, and provided food and water *ad libitum*. Unless otherwise stated, an equal number of both male and female were included in each group for analyses. All animal procedures were performed in accordance with National Institutes of Health (NIH) guidelines under protocols 2018-0215 and 2021-0292 approved by the Institutional Animal Care and Use Committee (IACUC) at Washington University School of Medicine in St. Louis, MO.

### Design of TPP1LAMP1 transgene and lentiviral transduction

The 114-bp transmembrane region and cytosolic tail of LAMP1 was linked in frame to the C terminus of 1689-bp human TPP1 cDNA (CCDS7770.1) via a six-glycine linker to retain TPP1 within the lumen of the lysosome. The Kozak sequence was added at the 5’ of TPP1 cDNA, and a 30-bp c-Myc epitope tag was linked to the cytoplasmic tail of the LAMP1 domain to complete the TPP1LAMP1 transgene. The TPP1LAMP1 transgene and the regular hTPP1 cDNA were each synthesized and inserted into the multiple cloning site of pLenti-III-PGK plasmid vector (Applied Biological Materials, Canada, Cat. No. G305). TPP1LAMP1 and TPP1 lentiviruses (LV-TPP1LAMP1 and LV-TPP1) from the plasmid were generated at the Hope Center Viral Vector Core at Washington University School of Medicine (St. Louis, MO). Immortalized *Cln2^−/−^* mouse embryonic cells (MEFs, kind gift from Dr. David Sleat) were transduced with either LV-TPP1LAMP1 or LV-TPP1. Selection was performed by adding 3 µg/ml puromycin to the media 96 hours post-transduction and continued for a week.

### Tissue culture

WT, *Cln2^−/−^*, and LV-TPP1 or LV-TPP1LAMP1 transduced *Cln2^−/−^* MEFs were cultured in DMEM High Glucose (Gibco, 19965) supplemented with 15% fetal bovine serum (Gibco, 10-437-028) and 1% penicillin-streptomycin (Sigma, P0781) on 6-well plates. When reaching 70-80% confluency, the media was replaced, and both cells and conditioned media were harvested 48 hours later for TPP1 activity assays. For the co-culture experiments, WT, *Cln2^−/−^*, and LV-TPP1 or LV-TPP1LAMP1 transduced *Cln2^−/−^* MEFs were seeded onto 6-well plates at approximately 20% confluency. After 24 hours, new *Cln2^−/−^* MEFs were placed on 24 mm Transwell inserts (Corning, 3450) and co-incubated with pre-seeded MEFs. After 72 hours, *Cln2^−/−^* MEFs from the inserts were harvested for the TPP1 activity assays.

### TPP1 enzyme activity assays

TPP1 enzyme activity in samples was measured using fluorometric assays, as described previously (24, 76). See Supplementary Methods for further details.

### Generation of TPP1LAMP1 transgenic mice

Using the pLenti-III-PGK-TPP1LAMP as a template. Poly-A tail signal was added at the downstream of c-Myc domain, and the sequence from the PGK promoter to the poly-A tail was flanked by *loxP* loci and mouse *Rosa26* homology arms, which was subsequently inserted into the inverted terminal repeats (ITRs) of the AAV2 vector (AAV2-TPP1LAMP1). Transgenic mice were generated by a combination of transduction of the AAV2-TPP1LAMP1 donor vector and electroporation of CRISPR Cas9 and the Rosa26-targeting guide RNA into *Cln2^R207X/WT^* embryos on the C57BL/6J background. Stable integration of the transgene was confirmed by PCR targeting both 5’ and 3’ junctions in the tail DNA. Fonder mice were crossed with *Cln2^R207X/R207X^* mice to obtain *TPP1LAMP1^flox/-^*; *Cln2^R207X/R207X^* mice (F1), and F1 mice were crossed with each other to obtain *TPP1LAMP1^flox/flox^*; *Cln2^R207X/R207X^* mice (= TPP1LAMP1 mice, F2). Homozygosity and heterozygosity of the TPP1LAMP1 allele was distinguished by PCR targeting the mRosa26 sequence outside the transgene insertion using the following primers. See Supplementary Methods for the information of PCR primers.

### CI-MPR affinity assays

CI-MPR assays were performed as previously described (29, 78). Briefly, soluble CI-MPR was purified from fetal bovine serum (FBS) and covalently conjugated to Cyanogen bromide-activated Sepharose 4B (Sigma-Aldrich). Total protein extracts from brains of WT and TPP1LAMP1 mice were diluted with buffer A (50 mM sodium acetate, pH 4.0, and 1% Triton X-100) and incubated with the CI-MPR beads at 4°C for 2 hours. After incubation, the beads were collected, washed twice with PBS and 1% Triton X-100, washed once with 0.1M sodium acetate buffer, and assayed for TPP1 activity. Affinities to CI-MPR are reported as the percentage of the starting enzyme recovered on the beads.

### Immunohistochemistry and imaging

Processing of forebrains were performed, as described previously (79–81). See Supplementary Methods for further details. A one-in-twelve series of 40µm coronal forebrain sections from each mouse were stained using on on-slide immunofluorescence protocol as previously described (82, 83). See Supplementary Methods for further details including primary and secondary antibodies. To visualize GABAergic interneurons for stereological counts, a one-in-six series of coronal sections were stained using a free-floating immunoperoxidase protocol (79, 81). See Supplementary Methods for further details including used antibodies. All images were taken on a Zeiss *AxioImager Z1* microscope with *StereoInvestigator* (MBF Bioscience) software or a Zeiss LSM880 Confocal Laser Scanning Microscope with *Airyscan* and *ZEN 2* (blue edition, Zeiss) software.

### Quantitative thresholding image analysis

To quantify storage material accumulation (SCMAS immunoreactivity) and glial activation (GFAP-positive astrocytes and CD68-positive microglia), semiautomated thresholding image analysis was performed as described previously (82, 83). This involved collecting slide-scanned images at 10x magnification (Zeiss Axio Scan Z1 Fluorescence Slide Scanner) from each animal. Contours of appropriate anatomical regions were then drawn, and images were subsequently analyzed using *Image-Pro Premier* (Media Cybernetics) using an appropriate threshold that selected the foreground immunoreactivity above the background. All thresholding data (GFAP, CD68 and SCMAS) were expressed as the percentage of area within each anatomically defined region of interest that contained immunoreactivity above the set threshold for that antigen (“% immunoreactivity”).

### Co-localization image analysis

To quantify the extent of overlap between GFAP (Alexa Fluor 488) and GABA (Alexa Flour 680) immunoreactivity, co-localization analysis was performed using *Zen 2* (blue edition, Zeiss) software on the confocal images taken with a 20x objective. Briefly, z-stack images covering the entire thickness of the section were processed using the maximum intensity projection function. The whole area of the images was analyzed using colocalization tools with appropriate thresholds set to distinguish foreground immunoreactivity from background for both channels. Three images were randomly selected per brain region per animal, and the average of the colocalization coefficients (area positive for both GFAP and GABA/area positive for GFAP) was reported as results.

### Stereological counts of interneuron number

Unbiased design-based optical fractionator counts of GABAergic interneurons were performed using *StereoInvestigator* (MBF Bioscience) in a one-in-six series of forebrain hemisections as described previously (79, 81, 82). Neurons expressing PV, SST, and CTIP2 with clearly identifiable immunoreactivity were counted using the optical fractionator method with the following sampling scheme and objective. PV and SST in S1BF, counting frame 180 × 200 μm, sampling grid 500 × 500 μm, objective 40×/0.95 Korr; CTIP2 in CPu, counting frame 100 × 10 μm, sampling grid 1000 × 1000 μm, objective 63× oil (NA 1.4).

### Total RNA extraction, cDNA synthesis, and PCR

Total RNA was extracted from liver homogenates from TPP1LAMP1 mice or HEK293 cells pellets (control), and purified using TRIzol (Thermo Fisher), as previously described (84). Subsequently, complementary DNA (cDNA) was synthesized using *Random Primers* (Invitrogen) and *SuperScript II* Reverse Transcriptase (Invitrogen) according to the manufacturer’s protocol. The presence of hTPP1 and TPP1LAMP1 transcripts was confirmed by PCR on cDNA using a forward primer targeting hTPP1 in conjunction with either a reverse primer targeting hTPP1 or a reverse primer targeting the LAMP1 domain. See Supplementary Methods for primer sequences.

### Quantitative gait analysis

The *CatWalk XT* gait analysis system (Noldus Information Technology, Wagenigen, Netherlands) was used to study gait performance at monthly intervals, as described previously (82, 85). See Supplementary Methods for further details.

### Electroencephalography (EEG) monitoring

Experimental mice underwent continuous video-EEG monitoring starting at 11 weeks, using standard methods for implanting epidural electrodes and EEG recording under isoflurane anesthesia, as previously described (86–88). Continuous bilateral cortical video-EEG signals starting at 11 weeks were acquired using a referential montage using *Stellate* or *LabChart* (AdInstruments) acquisition software and amplifiers until *Cln2^R207X^* mice died. See Supplementary Methods for further details.

### Seizure induction

Pentylenetetrazole (PTZ, Sigma) dissolved in phosphate-buffered saline (PBS) was injected intraperitoneally (75mg/kg) into 25-week-old TPP1LAMP1 or Vgat-Cre; TPP1LAMP1 mice, as previously described (89). Mice were then individually placed into cages for observation via video recordings until death. Generalized tonic-clonic seizure was characterized by sudden loss of upright with diffuse tonic posturing followed by clonic shaking. The minimal interval of 30 seconds between seizures was implemented for analysis.

### Chronic DREADD studies

Intracerebroventricular injections of 5 μL of either AAV9-hSyn-DIO-hM3Dq-mCherry (Addgene # 44361-AAV9) (90) or AAV9-hSyn-DIO-mCherry (Addgene # 50459-AAV9) vector at a concentration of 5 × 10^10^ gc per mouse were performed in P1 or P2 neonatal *Cln2^R207X/R207X^* mice. For EEG monitoring, administration of 10 µg/ml deschloroclozapine (DCZ, Hello Bio #HB9126) via drinking water was initiated from 11 weeks (78 days) of age. DCZ-containing drinking water was refreshed every three or four days. For histological studies, micropipette-guided DCZ administration, as previously described (41) was used. Briefly, animals at 11 weeks (78 days) of age were trained to drink 2 ml/kg of the 40% condensed milk solution dissolved in PBS from a single-channel P200 micropipette over three consecutive days, with one training session per day. From day 4 onwards, mice received daily doses of 500 µg/kg of DCZ dissolved in the same volume of 40% condensed milk. Mice were taken down 24 hours after the third DCZ administration (96 hours after the first DCZ administration) for histological analysis. This dose of DCZ was determined based on our pilot study involving WT mice expressing hM3Dq-DREADDs, where doses ranging from 1 to 100 µg/ml (in drinking water) and from 10 to 1000 µg/kg/dose (in condensed milk) were assessed for increased c-Fos expression as a marker of neuronal activation.

### Statistical analysis

All statistical analyses were performed using *GraphPad Prism* version 9.1.0 for MacOS (GraphPad Software, San Diego, CA). Unpaired t-test were used for comparison between two groups based on distributions of data. A one-way ANOVA with a post-hoc Bonferroni correction was used for comparison between three groups or more. Log-rank (Mantel-Cox) test was used for the onset of spontaneous seizures and survival studies. A p-value of ≤0.05 was considered significant.

## Date Availability Statement

All data discussed in the paper are either within the manuscript or will be made available to readers via contacting JDC directly.

## Acknowledgements

We acknowledge Dr. David Sleat (Rutgers New Jersey Medical School) for providing *Cln2^−/−^* and control WT MEFs that we used in our study. We thank Dr. Jill M. Weimer and Dr. David A. Pearce (Sanford Research) for originally supplying the *Cln2^R207X^* mice that were used in this study. We thank Dr. Takafumi Minamimoto (National Institutes for Quantum Science and Technology, Japan) for his expert advice on DCZ dosing. We also thank the Washington University Center for Cellular Imaging (WUCCI) supported by Washington University School of Medicine, the Children’s Discovery Institute of Washington University and St. Louis Children’s Hospital, and the Foundation for Barnes-Jewish Hospital in St. Louis, for providing a Zeiss Axio Scan Z1 Fluorescence Slide Scanner, the Molecular Microbiology Imaging Facility at Washington University School of Medicine in St. Louis for providing a Zeiss LSM880 Confocal Laser Scanning Microscope with *Airyscan,* the Hope Center Viral Vector Core at Washington University School of Medicine in St. Louis, for generating lentivirus, and the Genome Engineering & Stem Cell Center (GESC) and the Mouse Genetics Core (MGC) at Washington University School of Medicine in St. Louis, for generating TPP1LAMP1 transgenic mice. We acknowledge KT’s thesis committee members Drs. Joseph Dougherty, Gilbert Gallardo, and Tristen Qingyun Li, Washington University in St. Louis for their scientific input. We also acknowledge Dr. Alison Barnwell for constructive comments on the manuscript.

---

**Figure S1.**
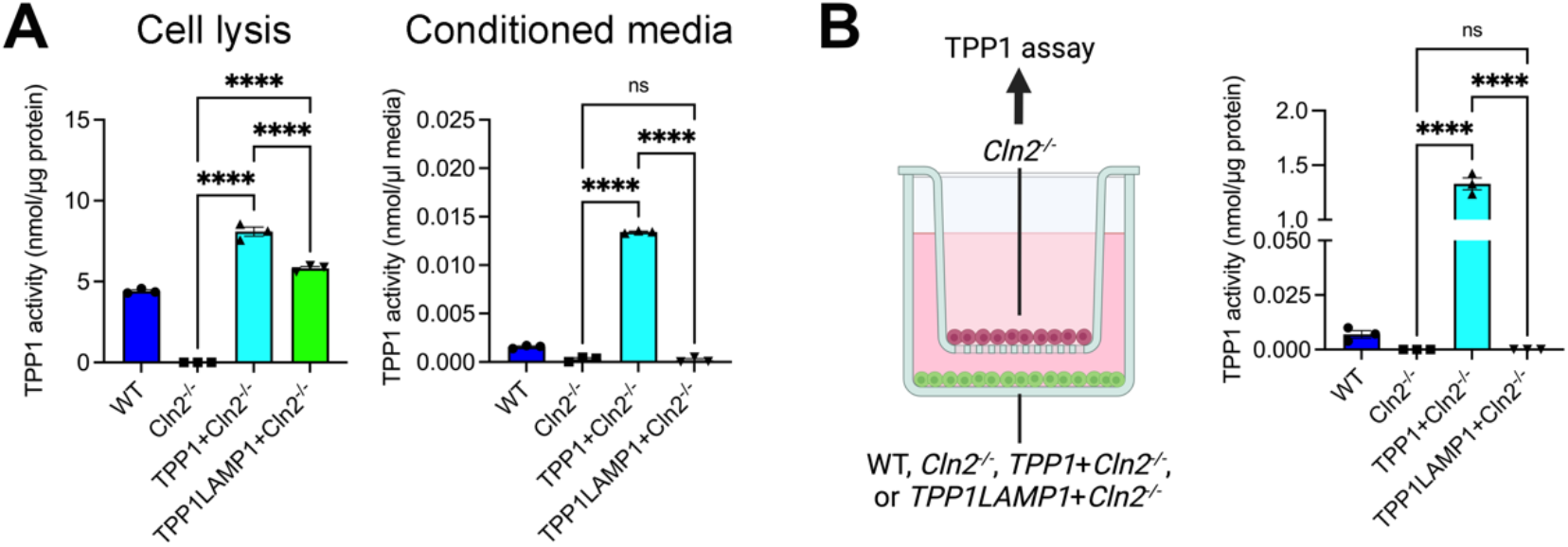
*In vitro* validation of TPP1LAMP1 construct. (*A*) TPP1 activity assay in both cell lysis (left) and conditioned media (right) of WT, *Cln2^−/−^*, *Cln2^−/−^*; TPP1-expressing *Cln2^−/−^*, and TPP1LAMP1-expressing *Cln2^−/−^* mouse embryonic fibroblasts (MEFs) shows intact intracellular activity with no extracellular secretion of TPP1LAMP1 *in vitro*. (*B*) Schematic describing the co-culture experiments using cell-inserts (left). The image was created with BioRender.com. TPP1 activity assay in *Cln2^−/−^* MEFs shows detectable TPP1 activity after co-cultured with WT and TPP1-expressing *Cln2^−/−^*, but no TPP1 activity after co-cultured with TPP1LAMP1-expressing *Cln2^−/−^* MEFs (right). Values are shown as mean ± SEM (n = 3 replicates per group). one-way ANOVA with Bonferroni correction. \**P* < 0.05, \*\**P* < 0.01, \*\*\**P* < 0.001, \*\*\*\**P* < 0.0001.

**Figure S2.**
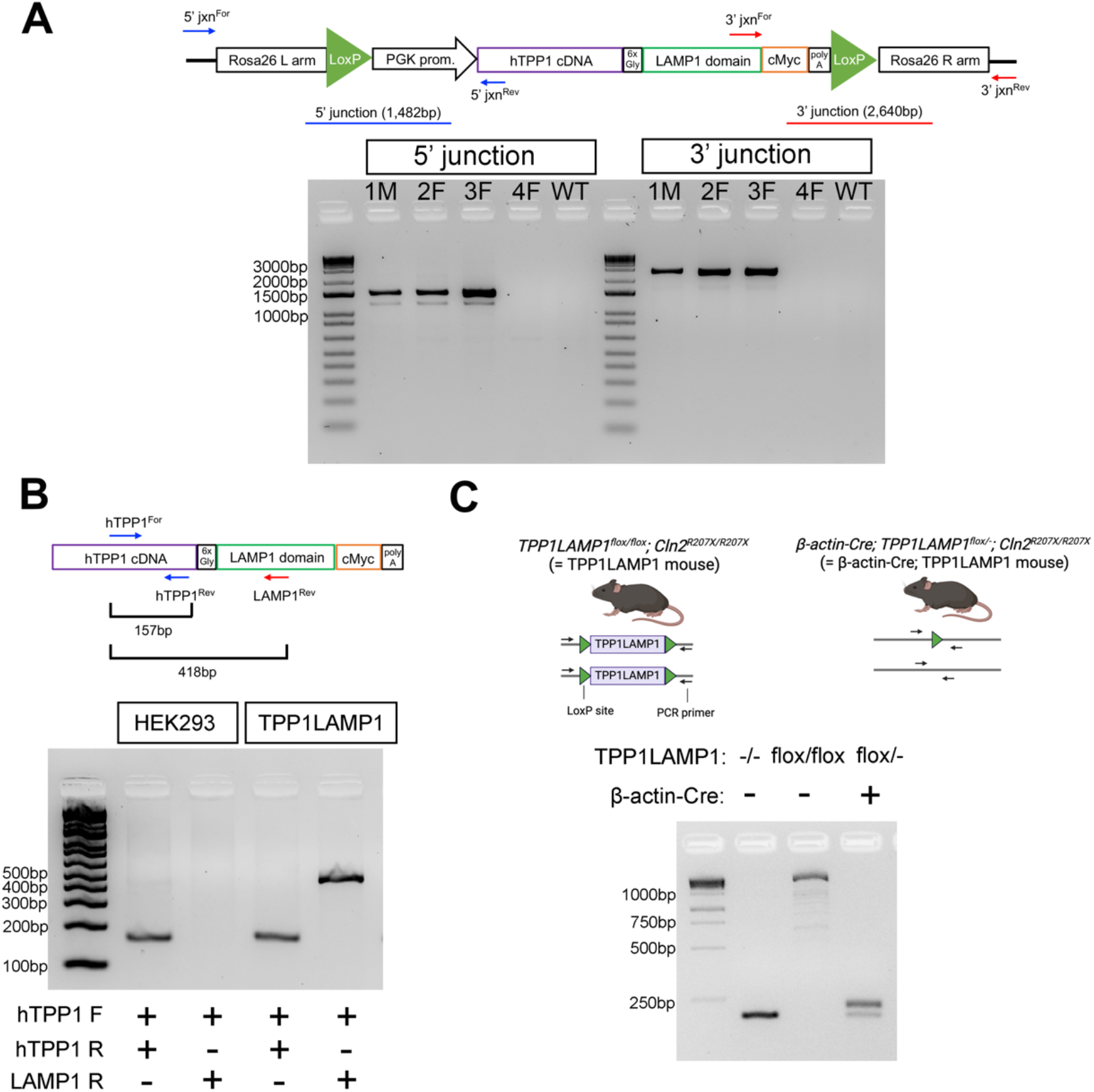
Validation of TPP1LAMP1 insertion and transcripts. (*A*) PCR targeting 5’ and 3’ junction of the TPP1LAMP1 insert at the Rosa26 locus confirms the intact insertion of the transgene. (*B*) PCR analysis on complementary DNA derived from mRNA extracted from the liver of TPP1LAMP1 mice (lane 3 and 4) shows the presence of the TPP1LAMP1 transcript at 418 bp. Control cDNA derived from HEK293 cells (lane 1 and 2) only shows the presence of hTPP1 at 157 bp. (*C*) Schematic describing the Rosa26 alleles in both TPP1LAMP1 mice and β-actin-Cre; TPP1LAMP1 mice (above). The images were generated with BioRender.com. PCR analysis targeting the entire *loxP*-flanked TPP1LAMP1 insert at the Rosa26 locus (below). WT mouse DNA (lane 1) shows homozygous WT alleles at 203 bp. TPP1LAMP1 mouse DNA (lane 2) shows homozygous TPP1LAMP1 alleles at 2898 bp. β-actin-Cre; TPP1LAMP1 mouse DNA (lane 3) shows the emergence of the floxed-TPP1LAMP1 allele at 258 bp on top of the WT allele at 203 bp.

**Figure S3.**
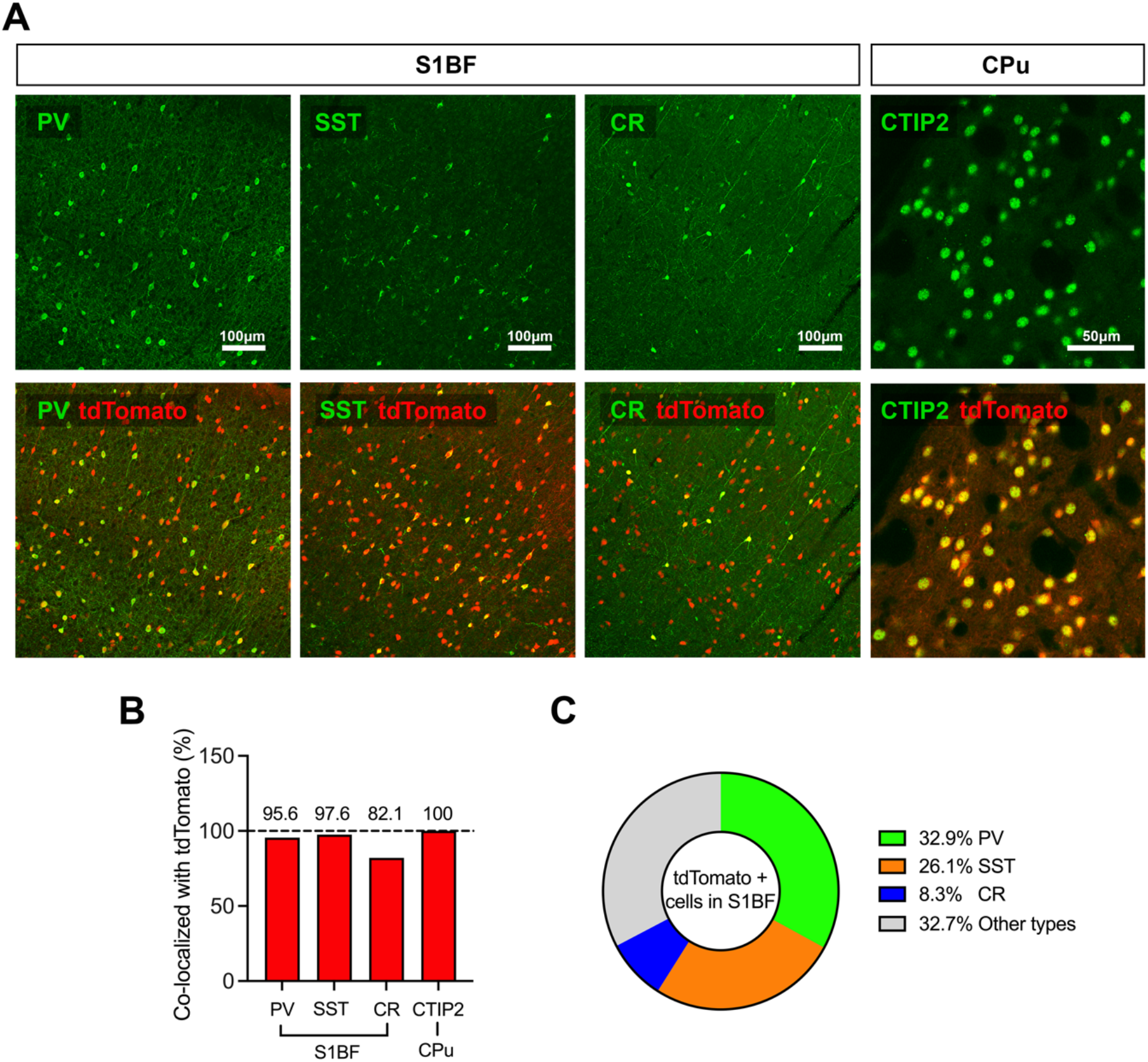
Validation of *Cre* expression pattern in *Vgat-Cre* mice. (*A*) Immunostaining for PV, SST, and CR (green) within S1BF and CTIP2 (green) within CPu shows these interneuron markers overlap with the endogenous tdTomato signal (red) in *Vgat-Cre^+/-^; Ai14^+/-^* mice signal at 8 weeks of age. (*B*) High percentages of PV-, SST-, and CR-positive interneurons within S1BF and CTIP2-positive interneurons within the CPu are positive for tdTomato in *Vgat-Cre^+/-^; Ai14^+/-^* mice. n=3. (*C*) Majority of tdTomato-positive cells within S1BF are PV- and SST-interneuron populations in *Vgat-Cre^+/-^; Ai14^+/-^* mice. n=3.

**Figure S4.**
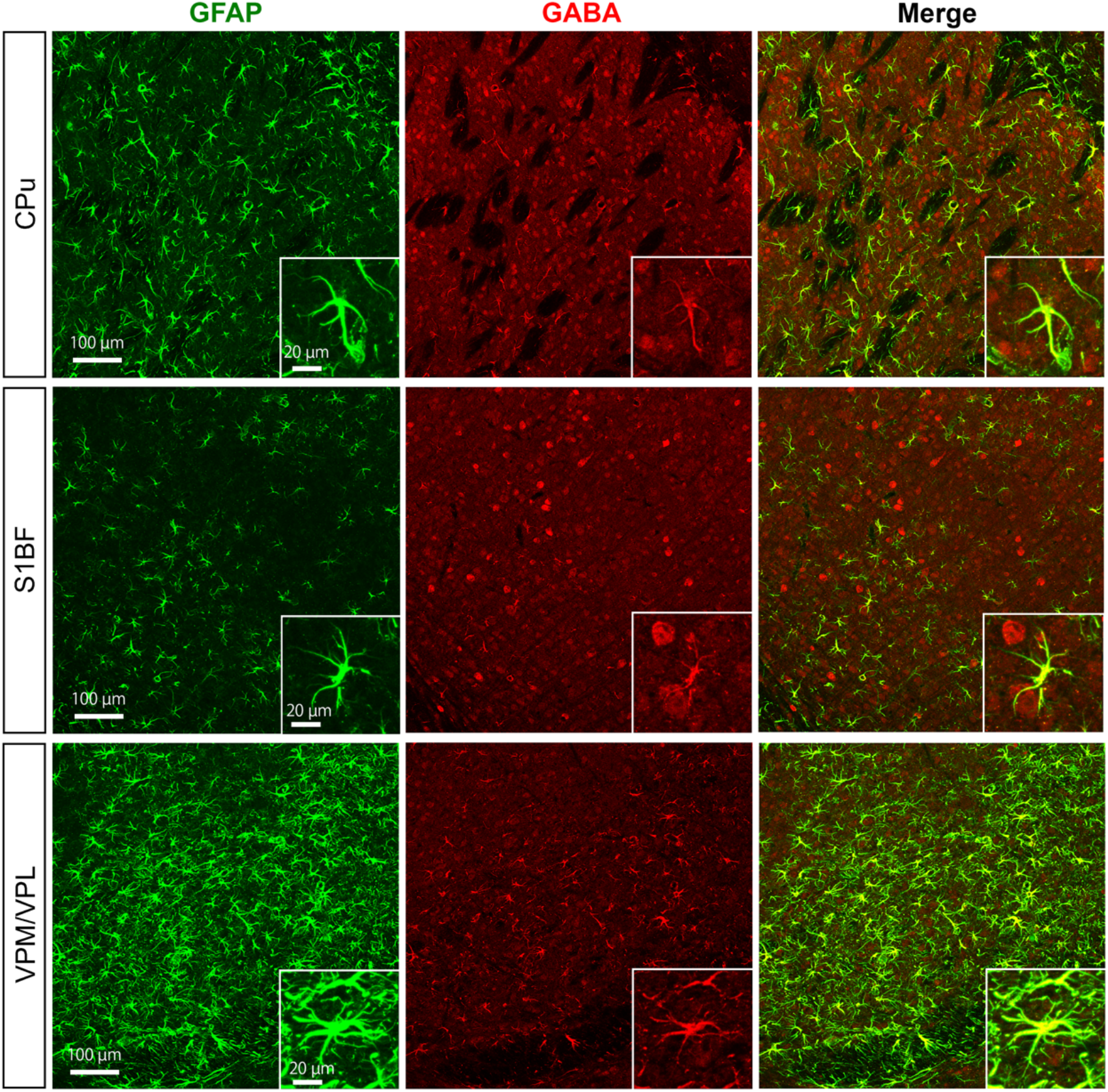
Increased GABA within astrocytes in *Cln2^R207X/R207X^* mice. Co-immunostaining for GFAP (green) and GABA (red) shows overlap between the two channels across CPu, S1BF, and VPM/VPL in 12-week-old *Cln2^R207X/R207X^* mice. Insets are higher magnification views from each image.

