## Supplementary Methods for "GABAergic interneurons contribute to the fatal seizure phenotype of CLN2 disease mice"

**Supporting Information**

**Detailed Materials and Methods**

*Tissue harvesting*

Mice were anesthetized using 2% isoflurane anesthesia and transcardially perfused with PBS. The brain was dissected out and bisected in the sagittal plane. One brain hemisphere was frozen and stored at -80°C for biochemical and molecular analysis, while the other brain hemisphere was fixed in 4% paraformaldehyde (PFA) solution for 48 hours and subsequently cryoprotected in 30% sucrose in 50 mM TBS (pH=7.6). Cryoprotected brain tissues were sectioned along the coronal axis at 40µm using a Microm HM430 freezing microtome (Microm International, Germany) equipped with a Physitemp BFS-40MPA freezing stage (Physitemp, Clifton, NJ) prior to histological analysis. Serum was separated from whole blood transcardially obtained from mice in regular 1.5ml microtubes, followed by centrifugation at 5000 rpm at room temperature. The resulting supernatant was collected and stored at -80°C for TPP1 enzyme activity assays.

*TPP1 enzyme activity assays*

TPP1 enzymatic activity in samples was performed using fluorometric assays, as described previously (1, 2). Briefly, upon removal from -80˚C storage, frozen forebrain tissues or cell lysates were homogenized in lysis buffer (50 mM sodium acetate pH 4.0; 0.1% Triton X-100). Homogenates were centrifuged at 12,000g for 25 min at 4˚C. Total protein concentration in each supernatant was measured using the Pierce BCA Protein Assay (Thermo Fisher Scientific), using a 96-well plate format. 10 μg of total protein extracts or 50 µl of tissue culture conditioned media were loaded into each well with 100 μl final volume of 100 μM Ala-Ala-Phe AMC (Sigma Aldrich) diluted in substrate buffer (0.1M sodium acetate pH 4.0; 0.1% Triton X-100) and incubated at 37˚C for 1 hour (brain) or 12 hours (cell lysis or conditioned media) and stopped by adding 50 ml of 0.4M glycine carbonate buffer (pH 10.8). TPP1 enzymatic activity in serum from mice was also measured by the same method, however due to limited volume of serum, 10 µl of serum was added into 50 µl of 200 μM Ala-Ala-Phe AMC (Sigma Aldrich) diluted in substrate buffer and incubated for 2 hours. Fluorescence intensity was measured in a Synergy H1 microplate reader (BioTek; Excitation 380 nm/Emission 460 nm). Each assay was done in triplicates with the average values used for analysis.

*Primers for PCR*

Genotyping for TPP1LAMP1 mice

5’ junction: forward, 5’-CGTCTCGTCGCTGATTGGCTT-3’, reverse, 5’-CAAAGAGCCCTAGGAGGCAGG-3’.

3’ junction: forward, 5’-CCAACTTCCCAGCTTTGCTGAAG-3’, reverse, 5’-CCCACAGTGTCAAAAGACCCAACC-3’.

mRosa26: forward, 5’-GAGTTCTCTGCTGCCTCCTG-3’, reverse, 5’- TGGGAAGTCTTGTCCCTCCA-3’.

PCR for complimentary DNA

hTPP1 forward, 5’-TTCTGATGGCTACTGGGTGG-3’

hTPP1 reverse, 5’-GAGCCTTGGGTTGAGAAAGC-3’

LAMP1 reverse, 5’-GAGGTAGGCAATGAGGACGA-3’

*Immunohistochemistry*

Immunofluorescence protocol: a one-in-twelve series of coronal forebrain sections were mounted on 25 x 75 mm microscope slides (Fisher Scientific) and air-dried for 30 minutes before blocking in Tris-buffered saline (TBS) with 4% Triton X-100 (Alfa Aesar) and 15% normal goat serum (Vector) for 1 hour at room temperature. Sections were then incubated for the following primary antibodies in TBS with 4% Triton X-100 and 10% normal goat serum for 2 hours at room temperature: rabbit anti-GFAP 1:1000, DAKO #Z0334, mouse anti-GFAP 1:500, Sigma #G3893, rat anti-CD68 1:400, Bio-Rad #MCA1957, rabbit anti-SCMAS 1:400, abcam #ab181243, mouse anti-parvalbumin (PV) 1:500, SWant #PV235, mouse anti-somatostatin (SST) 1:500, Santa Cruz Biotechnology #sc-55565, rat anti-COUP TF1-interacting protein 2 (CTIP2) 1:200, abcam #ab18465, rabbit anti-GABA 1:500, Sigma #A2052. mouse anti-mCherry 1:400, abcam #ab125096, and rabbit anti-c-Fos 1:400, EMD Millipore #ABE457. After 3 times of washing with TBS, sections were incubated for the following secondary antibodies in TBS with 4% Triton X-100 and 10% normal goat serum for 2 hours: Alexa Fluor 488 goat anti-rabbit IgG 1:400, Invitrogen #A11008, Alexa Fluor 546 goat anti-mouse IgG 1:400, Invitrogen #A11003, Alexa Fluor 546 goat anti-rat IgG 1:400, Invitrogen #A11081, and Alexa Fluor 647 goat anti-rabbit IgG 1:400, Invitrogen #A21244. If needed, sections were then incubated for NeuroTrace 640/600 Deep-Red Fluorescent Nissl Stain 1:100 in TBS for 20 minutes. After 3 times of washing with TBS, sections were quenched in 70% ethanol with 1x TrueBlack Lipofuscin Autofluorescence Quencher (Biotium) for 5 minutes, followed by 2 times washing with TBS and coverslipping with DAPI Fluoromount-G (SouthernBiotech).

Immunoperoxidase protocol: a one-in-six series of free-floating coronal forebrain sections were immunohistochemically stained using the standard immunoperoxidase protocol for interneuron markers. Briefly, sections were incubated in 1% H2O2 in TBS for 10 minutes at room temperature, rinsed in TBS three times, and blocked for 1 hour in TBS with 0.3% Triton X-100 and 15% normal goat or rabbit serum at room temperature before overnight incubation for the following primary antibodies in TBS with 0.3% Triton X-100 and 10% normal goat serum at 4˚C: rabbit anti-PV, 1:20000, SWant #PV27a, rabbit anti-Somatostatin (SOM) 1:2000, Immunostar #20067, and rat anti-COUP TF1-interacting protein 2 (CTIP2) 1:1000, abcam ab18465. After 3 times of washing with TBS, free-floating sections were incubated in biotinylated goat anti-rabbit secondary antibody (1:1000, Vector #VA1000) or biotinylated rabbit anti-rat secondary antibody (1:1000, Vector #BA4001) in TBS with 0.3% Triton X-100 and 15% normal goat or rabbit serum for 2 hours at room temperature, followed by 2-hour incubation in Vectastain ABC 1:1000 (avidin-biotin, 1:200, Vector) in TBS with 0.3% Triton X-100 at room temperature. Immunoreactivity was visualized using ImmPact DAB substrate kit, peroxidase (15 µl/ml, Vector #SK-4105) followed by airdrying for 2 hours and coverslipping with DPX mounting medium (Electron Microscopy Sciences).

*Quantitative gait analysis*

Gait analysis was performed using the *CatWalk XT* (Noldus Information Technology bv, Wagenigen, Netherlands) semi-automated gait analysis system, as described previously (3, 4). Briefly, mice were trained at least 2 days prior to data analysis and habituated to the room where behavior was performed overnight before testing. Each mouse needed to have run durations ranging from 1 to 5 seconds to be considered compliant. *Catwalk XT* 10.5 software (Noldus Information Technology) was then used to analyze different parameters associated with gait for each individual paw - Right Fore, Left Fore, Right Hind or Left Hind, in addition to removing any unwanted traces due to other parts of the body (e.g. mouse’s belly, scrotum or tail). Data for individual limbs were then averaged across all four paws in each run for individual animals where average data are shown. These data were then exported into an MS Excel spreadsheet (Microsoft, Seattle, WA).

*Electroencephalography monitoring*

Experimental mice underwent continuous video-EEG monitoring starting at 11 weeks, using standard methods for implanting epidural electrodes and EEG recording under isoflurane anesthesia, as previously described (5–7). Burr holes for the frontal reference electrodes were made (anterior +0.5mm, lateral +/-0.5mm; bregma) and secured via screws. Two bilateral “active” recording electrodes were placed over the parietal cortex (posterior -2.5mm, lateral +/- 1.5; bregma) and a ground screw secured over the cerebellum (posterior -6.2mm, lateral +/- 0.5; bregma). At least 72 hours after recovering from surgery groups of four mice were placed in individual cages and connected via a custom flexible cable attached to the exposed pin header for recording. Continuous bilateral cortical video-EEG signals starting at 11 weeks were acquired using a referential montage using Stellate or LabChart (AdInstruments) acquisition software and amplifiers until *Cln2^R207X^* mice died. Signals were amplified at 10,000X with high-pass (0.5Hz) and low-pass (100Hz) filters applied. EEG signals were digitized at 250 Hz and time-locked video EEG was collected continuously. Electrographic seizures were identified by their characteristic pattern of discrete periods of rhythmic spike discharges that evolved in frequency and amplitude lasting at least 10 seconds, typically ending with repetitive burst discharges and voltage suppression. Interictal epileptiform spikes and bursts of spikes were defined as a single sharp/fast (<200 ms) discharge or a burst of sharp discharges that disrupted the typical electrographic background and were greater than 2.5x amplitude compared to the surrounding background rhythm.

.
